# Self-transfecting GMO-PMO and PMO-GMO chimeras enable gene silencing *in vitro and in vivo* zebrafish model and NANOG Inhibition Induce the Apoptosis in Breast and Prostate Cancer Cells

**DOI:** 10.1101/2021.06.04.447039

**Authors:** Jayanta Kundu, Ujjal Das, Chandra Bose, Jhuma Bhadra, Surajit Sinha

**Affiliations:** School of Applied and Interdisciplinary Sciences, Indian Association for the Cultivation of Science, 2A and 2B Raja S. C. Mullick Road, Jadavpur, Kolkata 700032, India

**Keywords:** Morpholino Chimera, Self-transfecting antisense, Gene silencing, Hedgehog signaling, Zebrafish, NANOG, Cancer cells, Apoptosis

## Abstract

Phosphorodiamidate Morpholino Oligonucleotides (PMOs)-based antisense reagents cannot enter inside cells by itself without the help of any delivery technique which is the last hurdle for their clinical applications. To overcome this limitation, a self-transfecting GMO-PMO or PMO-GMO chimeras has been explored as a gene silencing reagent where GMO stands for guanidinium morpholino oligonucleotides which linked either at the OH- or NH-end of PMOs. GMO not only facilitates cellular internalization of such chimeras but also participates in Watson-Crick base pairing during gene silencing in ShhL2 cells when designed against m*Gli1* and compared with scrambled GMO-PMO where mutations were made only to the GMO part. GMO-PMO-mediated knockdown of *no tail* gene resulted no tail-dependent phenotypes in zebrafish and worked even after the delivery at 16-, 32- and 64-cell stages which were previously unachievable by regular PMO. Furthermore, GMO-PMO chimeras has shown the inhibition of *NANOG*, a key regulator of self-renewal and pluripotency of both embryonic and cancer stem cells. Its inhibition influences on the expression of other cancer related proteins and the respective phenotypes in breast cancer cells and increases the therapeutic potential of taxol. To the best of our knowledge, this is the first report on the self-transfecting antisense reagents since the discovery of guanidinium linked DNA (DNG) and most effective among the all cell-penetrating PMOs reported till date expected to solve the longstanding problem of PMO delivery. In principle, this technology could be useful for the inhibition of any target gene without using any delivery vehicle and should have applications in the fields of antisense therapy, diagnostic and nanotechnology area.

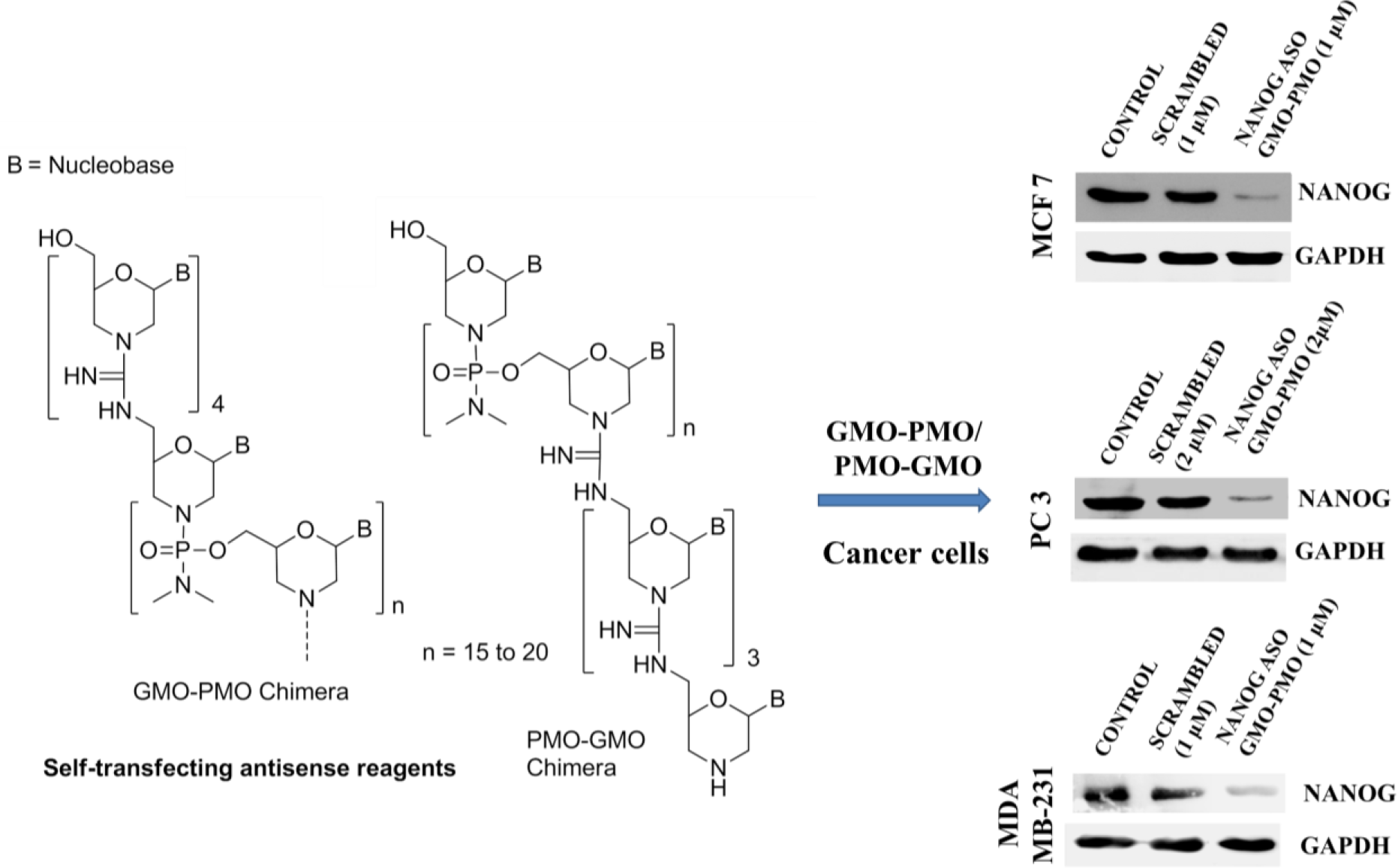

## Introduction

Phosphorodiamidate morpholino oligonucleotides (PMOs, **Figure 1a**) are unique class of antisense oligonucleotides because their ribose ring is replaced by morpholine ring and anionic phosphate backbone is replaced by neutral phosphorodiamidate backbone [1,2]. PMOs are routinely used as gene silencing reagents. PMOs bind to mRNA and block translation by a steric blockade mechanism when they are designed to be complementary to the 5′ leader sequences or to the first 25 bases 3′ to the AUG translational start site [1, 3]. Notably, two PMO-based drugs Eteplirsen [4] and Golodirsen [5] have been approved by the FDA for Duchenne muscular dystrophy (DMD), which is a hallmark for PMO-based antisense therapy that demonstrates the potential safety and effectiveness of PMO technology, is driving future development of such therapies for treatment of other diseases. However, similar to other oligonucleotides [6], the delivery of PMO is the long standing problem which is the main bottleneck for PMO’s clinical application [7]. At present guanidinium-rich cellular transporters are used for PMO’s delivery via the conjugation of PMO either with cell-penetrating peptides (CPPs) or cationic dendrimer to give PPMOs or Vivo-PMOs (**Figure 1a**) [8]. However, these oligos are associated with some level of toxicities due to the presence of too many cationic groups and the correlation between transfection efficiency and cytotoxicity of such transporters has been unsatisfactory [9‒11]. CPPs are peptide-based transporter and sensitive to peptidases. Modified version of CPPs have been reported for PMOs delivery in improving the bioavailability [12,13,14]. Generally, 8 to 12 cationic guanidinium groups are required for cell transfection, and hence prone to have non-specific interaction with negatively charged biomolecules, leading to toxicity.

**Figure 1.**
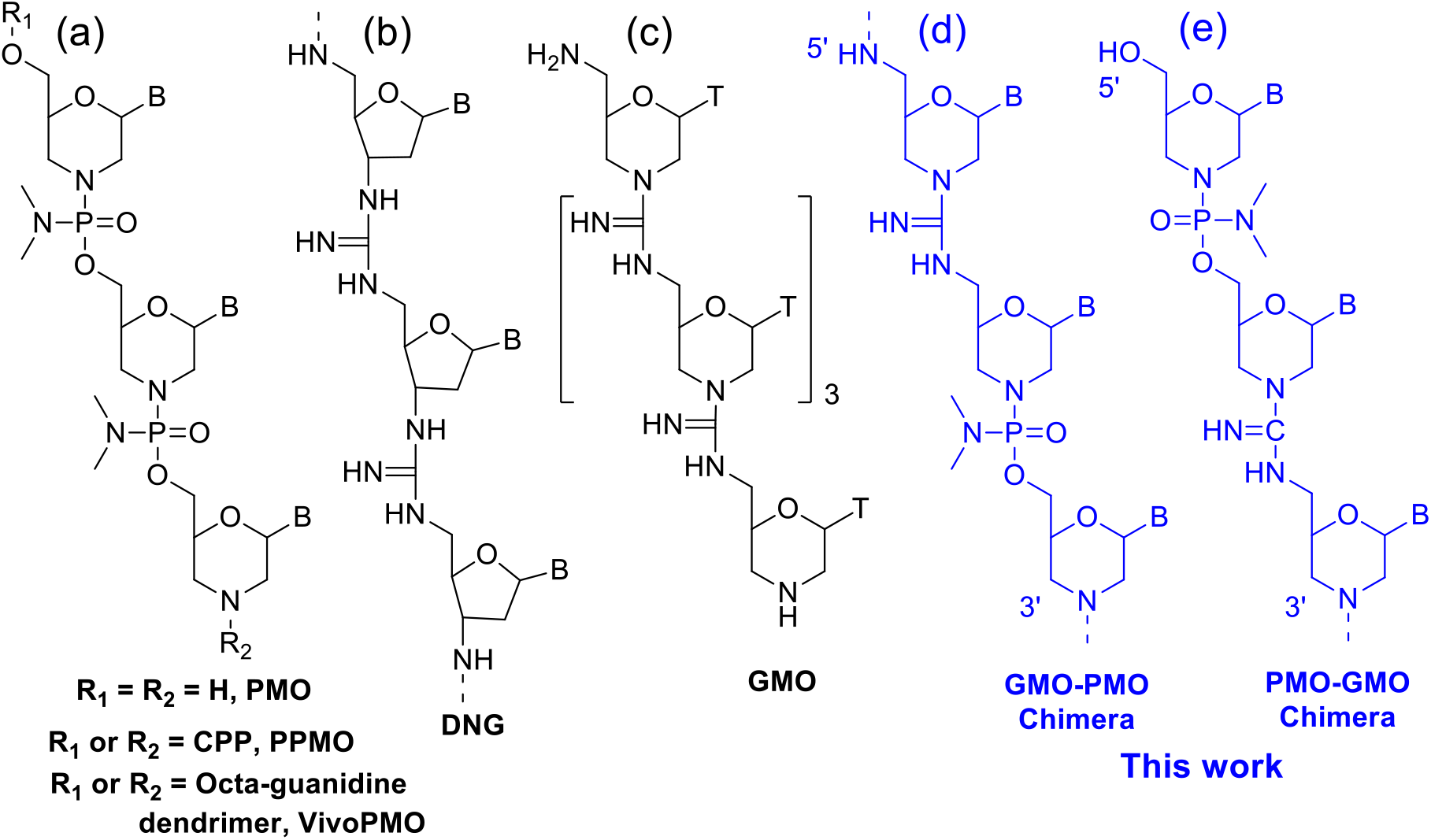
Chemical structures of (a) PMO, PPMO and VivoPMO; (b) DNG; (c) GMO; (d) GMO-PMO and (e) PMO-GMO chimeras.

To overcome the shortcoming associated with PMO delivery, we have developed different types of non-peptidic disubstituted internal guanidinium delivery moieties which successfully worked in PMO delivery with as minimum as four guanidinium linkages and was superior than vivoPMO concerning both efficacy and toxicity [15, 16].

With the continuous development of PMO delivery, our goal is to modify PMO in such a way that it will have effective antisense properties and at the same time it would be cell-permeable without any help of transporter or transfection reagents. As guanidinium rich molecules have a role for cell-penetration and guanidinium linked DNA analogues (DNG) (**Figure 1b**) have been reported by T. C. Bruice *et. al.* and DNGs are nuclease resistant and have high binding affinity towards DNA [17,18], however, to the best of our knowledge no gene silencing data is available for Bruice’s DNG till now except their cell-penetrating data recently reported by Mirkin *et. al.* [19]. While working on PMO, we initially introduced guanidinium linkages into morpholino-T nucleoside backbone to obtain guanidinium linked morpholino oligomer (GMO) (**Figure 1c**). Our initial penta-Thymidine GMO conjugated Gli1 PMO (25-mer) efficiently inhibit Gli1 expression level in Hedgehog signalling pathway [20]. We then became interested to incorporate the guanidinium linkages into PMO backbone to get at-a-stretch 20 to 25-mer GMO-PMO or PMO-GMO chimera which is expected to have both cell permeability and Watson Crick base pairing (**Figures 1d** and **e**). In such cases no further conjugation is required with PMO. Herein, we report the cell-penetrating and gene silencing properties of self-transfecting GMO-PMO and PMO-GMO chimeras against *Gli1* in Shh-Light 2 cells and *no tail (Ntl)* in zebrafish model and NANOG in breast cancer cells and NANOG-mediated apoptosis in cancer cells.

## Results and discussion

20-25-mer sequences of GMO-PMO and PMO-GMO chimeras (**Table 1**) were synthesized from respective activated monomers (data has not been published) in which PMO part was synthesized following our published protocol [21, 22].

**Table 1:**
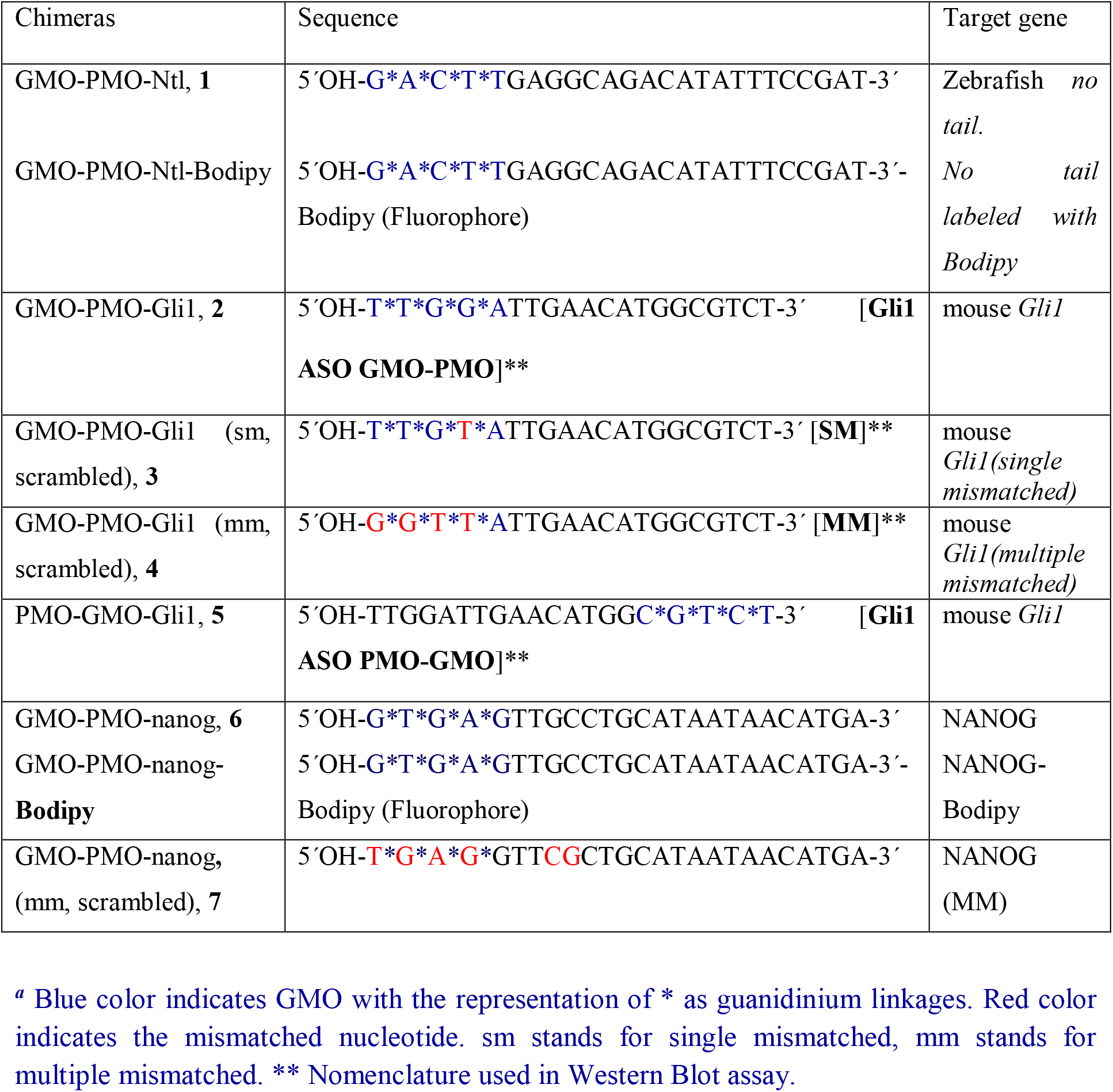
Sequences of GMO-PMO and PMO-GMO chimeras for gene silencing study.

In case of **GMO-PMO Ntl, Gli1** and **NANOG**, guanidinium linkages were incorporated at the 5′-end and in case of **PMO-GMO Gli1**, guanidinium linkages were incorporated at the 3′-end of the oligos. **GMO-PMO Ntl** was designed to target zebrafish *no tail* gene. **GMO-PMOs Gli1** and **PMO-GMO Gli1** have been designed targeting *Gli1*, including scrambled sequence (for control study). Gli1 is a transcription factor required for hedgehog signaling.

### GMO-PMO mediates suppression of *No tail* (*Ntl*) in zebrafish (*Danio rerio*)

For the evaluation of antisense application, zebrafish was chosen as a model organism because PMO are routinely used for gene silencing in zebrafish model [23, 24] and we used GMOPMO Ntl chimera against *no tail* (*ntl*) gene because no tail-dependent phenotypes are visible and reproducible [23]. As per previous report [16] 9 ng (0.25 mM stock, 3‒4 nl) of morpholino was injected into zebrafish embryos at 8 cell, 16 cell (1.6 hpf), 32 cell (1.75 hph) and 64 cell stages (30‒33 embryos were injected in each set). In all the cases Zebrafish no tail mutant phenotype was observed with a complete loss of vacuolated notochord cells, posterior structures and U-shaped somites rather than normal V-shaped (**Figure 2**). Zebrafish no tail mutant phenotypes were compared with wild type embryo and imaged at 30 hpf under microscope. Regular PMO injected after 32-cell stage did not show any phenotypes which indicated that self-penetrating **GMO-PMO 1** is capable of gene silencing even after 4 to 8 cell stages. To visualize the distribution of GMO-PMO **1** into zebrafish embryo, **Bodipy-conjugated GMO-PMO-Ntl** was also synthesized and injected into zebrafish embryo likewise GMO-PMO**-Ntl**. Under fluorescence microscope **Bodipy-GMO-PMO** injected embryo showed a widespread green fluorescent which imply a ubiquitous distribution of chimera into *in vivo* zebrafish model (**Supporting Figure 1 A, B**).

**Figure 2.**
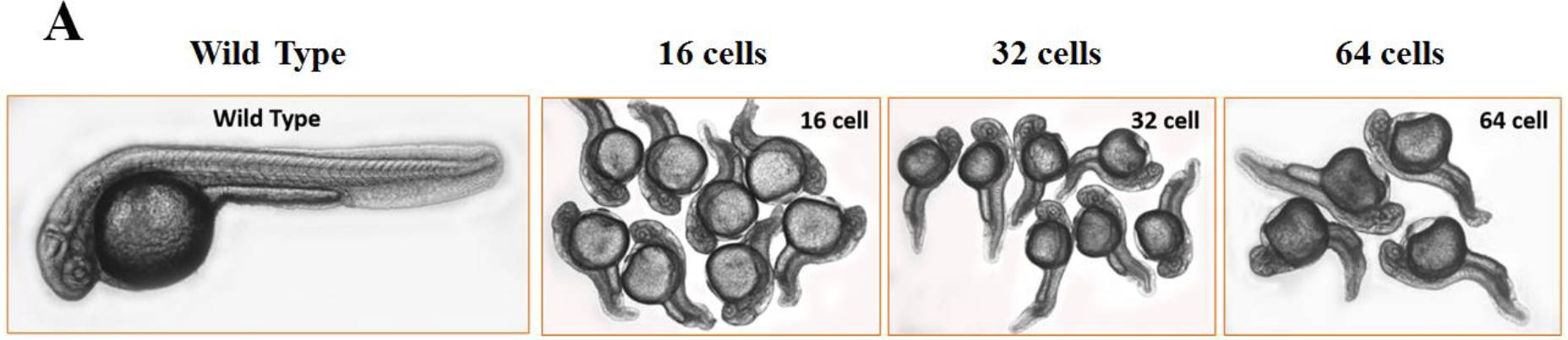
No tail GMO PMO injected zebrafish embryos in 16, 32 and 64 cell stages observed under phase contrast microscopes.

### Gli1 ASO GMO-PMO suppresses Gli1 and alters the expression of related genes

After this encouraging result, we then became interested to study the gene silencing efficacy of GMO-PMO chimeras in cell culture because it is notoriously difficult to deliver of PMO in cell culture model as PMO hardly form any complex with cationic lipids [25]. Though there are few methods of PMO delivery like electroporation [26], scrap loading [26] and endo-porter [27] or cellular transporter conjugated PMOs (VivoPMO [8], PPMO [8], IGT-PMO [15, 16], however, in the present case there is no need of using any delivery reagents or methods. In the attempt to provide the proof of principle we became interested in evaluating the antisense applications of GMO-PMO chimeras against Gli1, a transcription factor required for hedgehog signalling [28‒30]. Hh-Gli signalling is important during embryogenesis, stem cell differentiation and adult tissue homeostasis, whereas persistent Hh-pathway activity is implicated in the formation and growth of several cancers [29].

Next, we evaluated the antisense efficacy of GMO-PMO-Gli1 chimera in ShhL2 cells [derived from NIH3T3 (mouse embryonic fibroblast) cell line, stably transfected with Gli-dependent fire fly luciferase reporter] targeting Gli1 [31]. As the endogenous expression level of Gli1 is very low, the pathway was stimulated with Shh-conditioned media to obtain detectable amount of Gli1. Cytotoxicity assay by MTT was performed in ShhL2 cells using various doses of scrambled morpholino with four mismatched bases **GMO-PMO-Gli1 (MM) 4** and it was found to be nontoxic even as high as 70 μM dose which is much higher value than being used in the experiment (**Supporting Figure 2A**). In order to confirm the role of GMO part in gene silencing, we then used two 20-mer sequences (scrambled Morpholinos) with a single mismatched **GMO-PMO-Gli1 (SM) 3** and four mismatched sequences **GMO-PMO-Gli1 (MM) 4** in GMO part. Comparing with the normal 20-mer **GMO-PMO-Gli1 2**, there was no inhibition of Gli1 by scrambled morpholino with four mismatched bases **(MM), Figure 3A, B lane 3)** and about 40% inhibition was found in case of single mismatched **(SM) (Figure 3A, B, lane 2)**. It indicated that this GMO part is responsible for binding to mRNA. Moreover, immunoblot analysis also showed dose-dependent inhibition of Gli1 at 0.5 and 0.75 μM of Gli1 **ASO GMO PMO-Gli1**, which was not observed in case of **GMO-PMO-Gli1 (SM)** and **(MM)**(**Supporting Figure 2B, C**). In order to know the role of position of guanidinium linkages in PMO, **GMO-PMO-Gli1 2** and **PMO-GMO-Gli1 5** chimeras were synthesized with the incorporation of guanidinium linkage at the 5′- and 3′-end of PMO, respectively (data has not been published). Here we used both the Gli1 **GMO-PMO-Gli1** and **PMO-GMO-Gli1** chimeras. Efficacy wise, both the chimeras worked well whether guanidinium linkages are at the 5′-end or at the 3′-end (**Figure S 2D, E**). If we consider from synthetic point of view then initial incorporation of guanidinium linkages is better than incorporation at the 3′ end because it was easy to remove any truncated product if formed.

**Figure 3.**
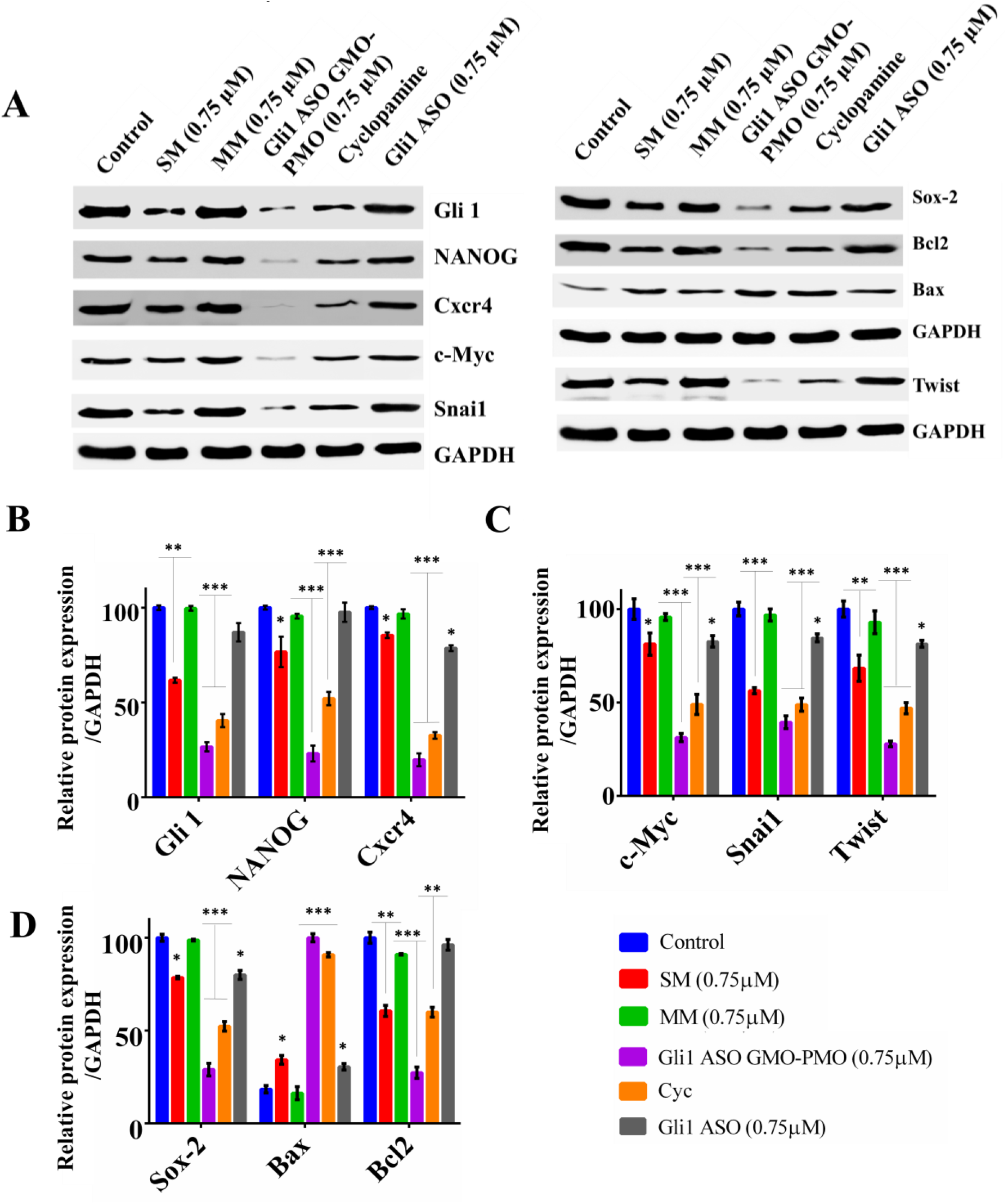
(A). Immunoblot data of pluripotency factors Nanog, Cxcr4, c-Myc, Sox2, EMT markers Snai1, Twist and apoptotic markers Bax, Bcl2 in Shh-Light2 cells after 36 hr of treatment. (B‒D). The densitometric analysis of immunoblot data. The protein expression levels are expressed relative to GAPDH. Error bars indicate means ± SE (n = 3), and data are presented as percentages relative to the control Shh Light 2 cells. *p< 0.05.

Next, we got intrigued to study the expression of other Hh related proteins in Shh Light 2 cells in order to observe the effect of this antisense treatment in downstream pathway. Like Gli1, the Gli1 **ASO GMO-PMO** treatment led to the down-regulation (approximately 75%) of NANOG and Cxcr4 at 0.75 μM dose which corroborated with the effect of Cyclopamine; a known inhibitor of Hh pathway (**Figure 3 A, B**). The suppression of Gli 1 by antisense oligo was also downregulated the embryonic stemness factors like c-Myc, Snai1, Twist, Sox-2 and anti-apoptotic protein Bcl2 in ShhL2 cells (**Figure 3 A-D**), indicating the significant involvement of Hh pathway in maintenance of proliferation, growth and self-renewal process of embryonic stem cells.

### GMO PMO Mediated NANOG Inhibition

Similarly the toxicity of GMO-PMO chimera was evaluated with scrambled ASO GMO-PMO. Significant decrease in cell viability was observed at a dose of 70 μM of scrambled oligo GMO PMO **7**(**Figure 4 A**) while the viability was significantly decreased even at 0.5 μM dose of NANOG **ASO GMO PMO 6**(**Figure 4 B**), indicating negligible toxicity of GMO PMO chimera at lower doses. Next, the antisense efficacy of GMO-PMO chimera **6** was evaluated in breast cancer cells (MCF7) where western blot analysis showed significant reduction of NANOG upto 52%, 76% and 82% inhibition at 0.5, 1 and 2 μM, respectively (**Figure 4 C, S3 A**). Since similar level of inhibition was obtained at 1 and 2 μM, hence the rest of the experiments were conducted with the lower dose. In our previous study, we observed that the NANOG **ASO IGT PMO**(5 μM) suppressed NANOG expression in MCF7 cells [15]. However, in present study when we compared the effect of NANOG **ASO GMO PMO 6** and NANOG **ASO IGT PMO**, better inhibition of NANOG was observed in case of NANOG **ASO GMO PMO 6** at 2 μM dose (**Figure 4 C, Figure S3 A**). Not only that, similar level of inhibitory effect on NANOG was seen after 36 and 72 hrs of incubation under both 0.5% and 10% serum concentrations (**Figure S3 B, C**) which indicated the presence of inter guanidinium linkages does not involve with serum interaction. It is an interesting observation like our previously reported IGT-PMO [**15**] as it does not happen in the case of other guanidinium-based cellular transporters [**32**].

**Figure 4:**
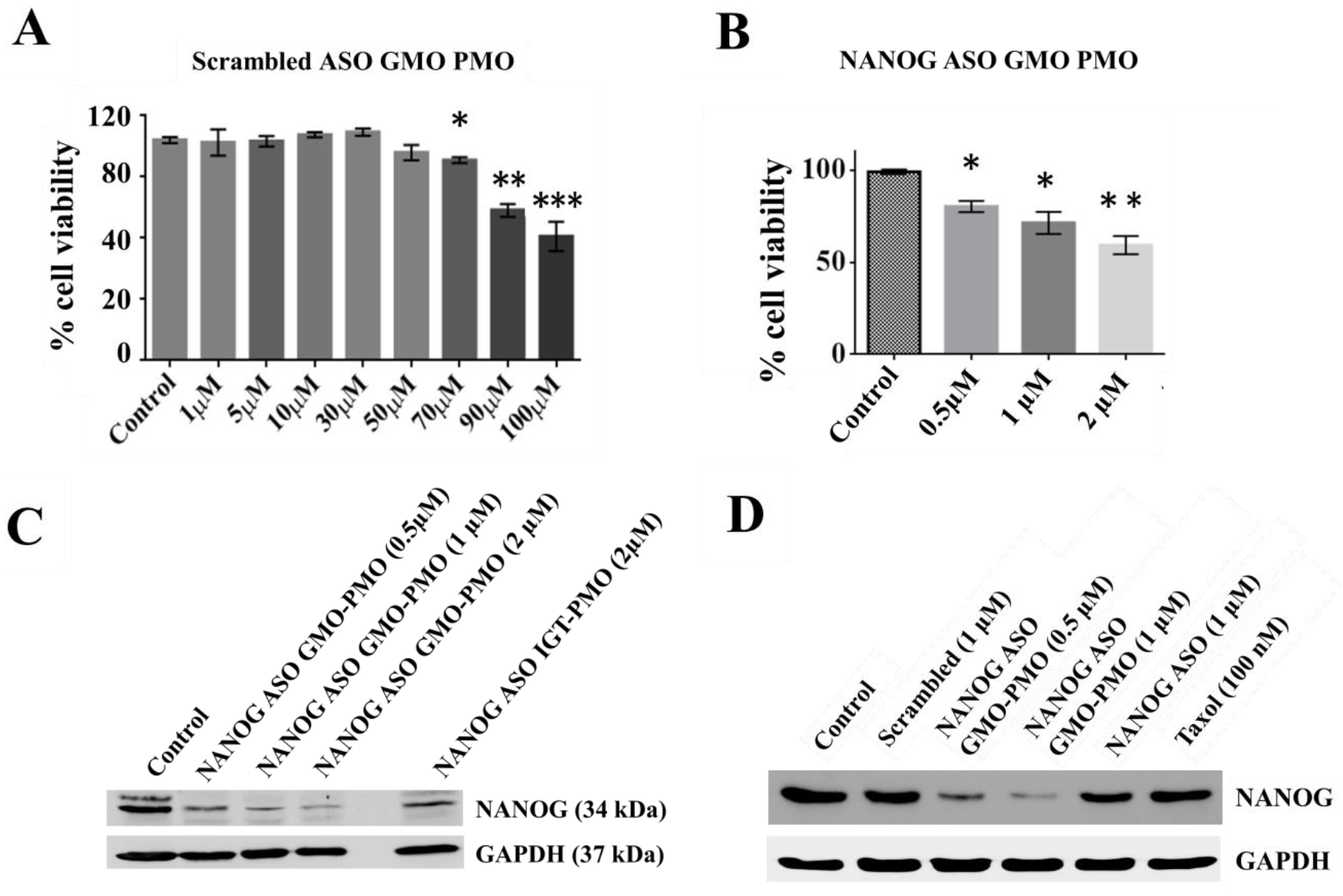
A. Cell viability assay by MTT and its graphical depiction in MCF7cells after treatment with different doses of scrambled NANOG oligo GMO PMO and B. with different doses of NANOG ASO GMO PMO. C. The immunoblot analysis of the expression of NANOG in MCF7cells at different doses of NANOG ASO GMO PMO w.r.t. 2μM of NANOG ASO IGT-PMO. D. Western blot analysis of the expression of NANOG in MCF7cells after treatment. NANOG ASO stands for naked PMO from Gene Tools.

### Inhibition of NANOG by GMO PMO abrogates the expression of stemness regulators

The pluripotency of the cancer stem cells (CSC) is determined by the concerted action of EpCAM, c-Myc, Sox2 and NANOG. For the directed migration and maintenance of these CSCs, NANOG cooperates with Cxcr4 playing a key role in perception of chemotactic gradients. On the other hand, phosphorylation Glycogen Synthase Kinase 3 beta (GSK3β) leads to the downregulation of stemness factors thereby inhibiting their maintenance and self-renewal. Thus it is clear that they are closely linked to each other and alteration of one affects the expression of others [33–36]. Likewise it was seen that inhibition of NANOG by GMO PMO led to the downregulation of Sox-2, cxcr4, EpCAM and c-Myc and upregulated the level of p-GSK3β (**Figure 4 D, Figure 5 A, B**).

**Figure 5:**
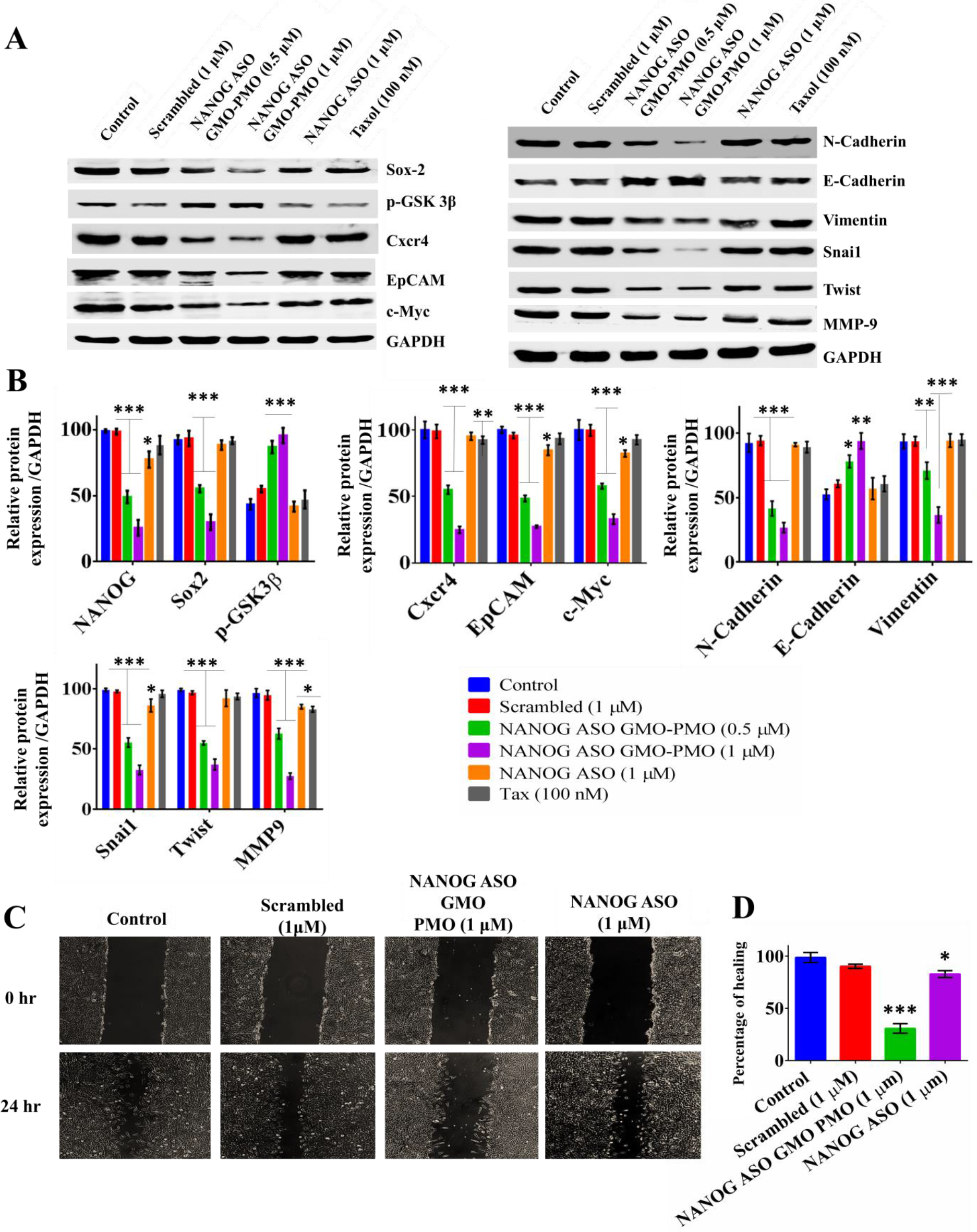
Nanog knockdown alleviates migration potential of MCF7 cells: A. NANOG ASO GMO PMO downregulates the expression of other related stemness factors such as Sox-2, p-GSK3β, Cxcr4, EpCAM, c-Myc and also other EMT markers such as N-Cadherin, Vimentin, Snai1, Twist, MMP9 as depicted from immunoblotting. B. The densitometic analysis of the immunoblot data. The protein expression levels are expressed relative to GAPDH. C. Wound healing assay in the cells upon indicated treatment where the width of wound closure in MCF7 cells at 0hr was set to 100%. D. Percentage of wound closure plotted graphically with regard to control. Scale bars represent 60 μm. Error bars indicate means ± SE (n = 3), and data are presented as percentages relative to the non-treated MCF-7 cells. *p< 0.05.

### Inhibition of Nanog by GMO PMO alleviates invasive and migratory potential of MCF7

Epithelial to Mesenchymal transition (EMT) plays a crucial role in tumour progression and metastasis in which epithelial cells lose cell polarity, cell-cell adhesion and gain mesenchymal properties such as migration and invasion properties [37]. EMT is also associated with attribution to stemness and resistance to radio and chemotherapy [38–41]. Thus, the expression of both epithelial and mesenchymal markers was evaluated after the treatment with NANOG **ASO GMO PMO 6**. Results indicated that after treatment with **6**, the expression levels of mesenchymal markers like N-cadherin and vimentin and other EMT proteins like Snai1, Twist and MMP9 decreased considerably with statistically significant upregulation in E-cadherin expression (**Figure 5 A, B**). This was also reflected in the wound healing assay which showed that the relative migration rates of NANOG inhibited cells significantly reduced to 65% compared to control. Thus, it is implicated that NANOG disruption by NANOG **ASO GMO PMO 6** has been effectual in preventing EMT in MCF7 cell line (**Figure 5 C, D**).

### Cellular uptake and endosomal escape study

Now, it is clearly understood that GMO-PMO chimeras can silence gene without any delivery vehicle or transfection reagents. We were then interested to see the cellular uptake, localization and distribution of NANOG **ASO GMO-PMO** chimera **6**. Accordingly, fluorescent version of **6** was synthesized by conjugating with Bodipy dye to obtain **6-Bodipy.** MCF7 cells were treated with **6-Bodipy** in time and dose dependent manner. The MCF7 cells were treated with 0.5 μM and 1 μM doses of bodipy tagged NANOG **ASO GMO-PMO** separately and the fluorescence was measured at different time points by flow cytometry. The incorporation of **6-Bodipy** was significantly increased with the increasing time of incubation after NANOG **ASO GMO-PMO** treatment and importantly 100% cell transfection was observed after 1 h of incubation (**Figure 6 A, B** and **S 4 A, B**). To identify the minimum concentration of **6-Bodipy** required to get 100% transfection, cells were treated with 50 nM to 500 nM of **6-Bodipy** for 4 h and 100% cell transfection was observed with 50 nM dose compared to control (**Figure 6 C** and **S 4 C**).

**Figure 6:**
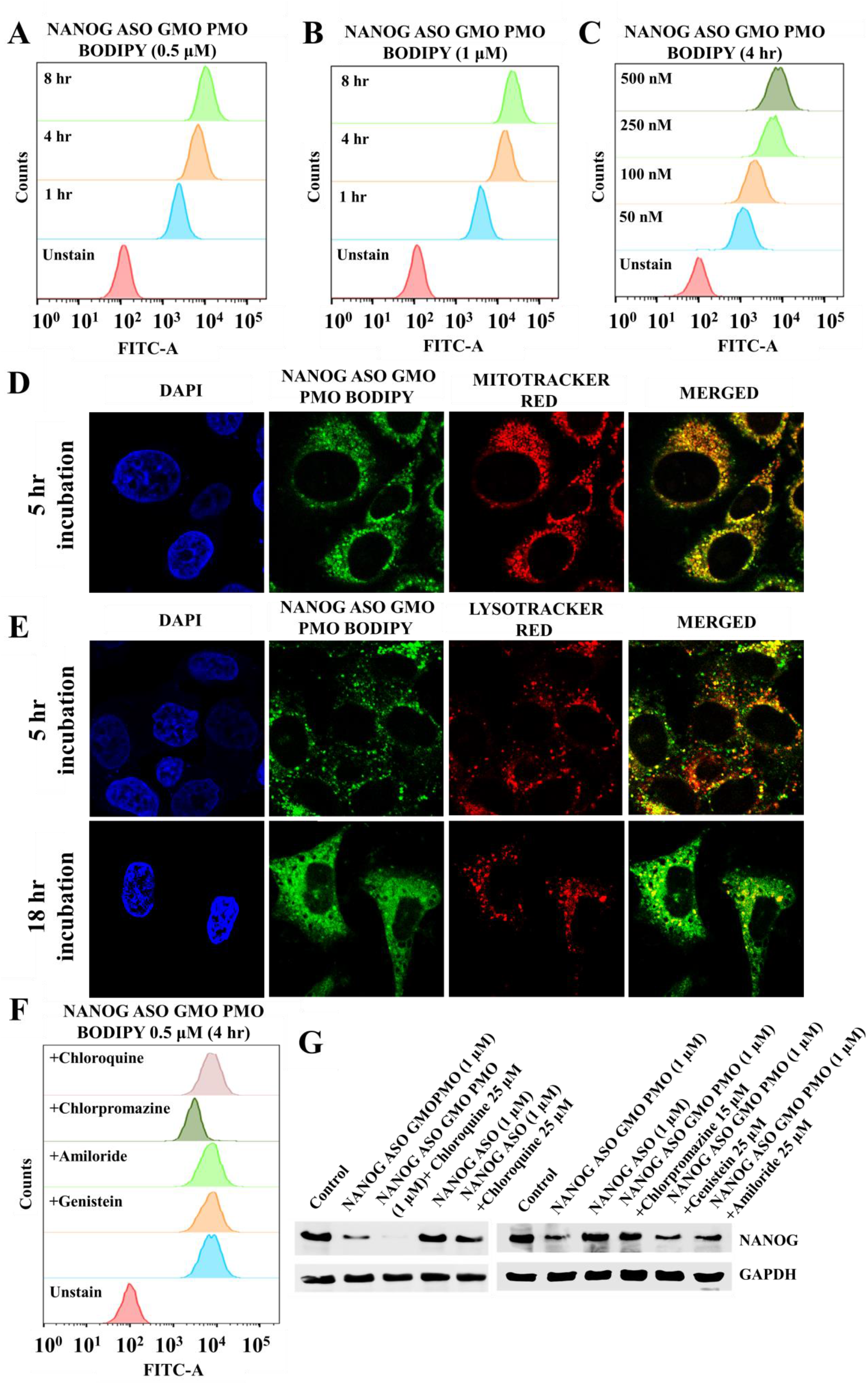
A. The overlapped histogram plots show the fluorescence intensity of bodipy NANOG ASO GMO-PMO with 0.5 μM dose B. 1 μM dose at different time points. C. The histograms show the dose dependent incorporation of antisense GMO-PMO in MCF-7 cells D. The confocal images show the localization of NANOG ASO GMO-PMO in mitochondria. Nuclei were stained by DAPI and Mitotracker red was used to stain mitochondria. E. The images show the localization of antisense oligo in lysosome and their endosomal escape. Lysotracker red was used to stain lysosome. F. The overlapped histograms show the fluorescence intensity of bodipy tagged GMO-PMO in presence of inhibitors of endocytosis pathway. G. Immunoblot analysis of the expression of NANOG in MCF7cells after treatment with NANOG ASO, NANOG ASO in combination with Chloroquine, NANOG ASO GMO PMO, and NANOG ASO GMO PMO in combination with Chloroquine, Chlorpromazine, Genistein and Ameloride. The protein expression levels are expressed relative to GAPDH. NANOG ASO stands for naked PMO from Gene Tools.

To understand the sub-cellular distribution of **6-Bodipy**, the cells were counterstained with Mitotracker red and Lysotracker red (50 nM) separately and observed under ultra-high resolution confocal microscope. After 5 h of incubation in 10% serum, bodipy tagged antisense oligo prominently co-localized with Mitotracker and Lysotracker red which therefore, proved their localization in mitochondria and lysosome at initial phase (**Figure 6 D, E** and **S 4 D, E**). Endosomal escape is a rate-limiting step to have a therapeutic efficacy of any drugs. In order to know the endosomal escape of chimera **6-Bodipy**(0.5 μM), the images were also captured after 18 h of incubation. Though the co-localization of NANOG **ASO GMO-PMO** was observed with lysosome even after 18 h, the major distribution of fluorescence was found in the cytosol which indicated that **6-Bodipy** was mostly escaped from endosomal compartment within 18 h of **6-Bodipy** treatment (**Figure 6 E** and **S 4 E**).

### NANOG ASO GMO PMO 6 enters through clathrin-mediated endocytosis pathway

There are three major endocytosis pathway operated in the MCF7 cells, clathrin, caveolin and macropinocytosis [42, 43]. In order to determine the mode of cytosolic delivery of **chimera 6**, the MCF7 cells were treated with **chimera 6** as well as with **6-Bodipy** separately with or without Chloroquine, Chlorpromazine, Genistein and Ameloride. The histogram plots of flow cytometry data showed significant increase of fluorescence intensity in MCF7 cells when treated with **6-Bodipy**. Moreover, no change in fluorescence intensity was observed when **6-Bodipy** was added to cells in combination with Chloroquine, Genistein and Amiloride. Surprisingly, treatment with Chlorpromazine prior to **6-Bodipy** significantly decreased the fluorescence intensity of **6-Bodipy** in MCF7 cells which proved the involvement of clathrin mediated endocytosis pathway for the entry of NANOG **ASO GMO-PMO**(**Figure 6 F, S 5 A**).

The MCF7 cells when treated with NANOG ASO alone, suppression of NANOG expression was observed in MCF7 but the level of inhibition was not sufficient to induce any morphological changes in MCF7 cells (**Figure 6 G** and **S 5 B, C, S6**). But pre-treatment with chloroquine increased the effect of NANOG ASO in terms of morphological alteration and NANOG suppression indicating enhanced endosomal escape (**Figure 6 G, S 5B**, **C, S6**). The similar effect of chloroquine was also found in case of NANOG **ASO GMO PMO**, where chloroquine pre-treatment increased the suppression of NANOG from 76% (NANOG ASO GMO PMO) to more than 93% (**Figure 6 G, S 5 B**). Chloroquine treatment didn’t increase the fluorescence intensity of **6-Bodipy** in MCF7 cells (**Figure 6 F, S 5 A**) rather it increased the endosomal escape and thus, the same concentration of NANOG **ASO GMO-PMO** showed better inhibition in presence of chloroquine. In order to confirm further the clathrin mediated endocytosis, we treated the cells individually with specific inhibitors of these pathways in presence of NANOG **ASO GMO PMO 6**. The genistein and amiloride did not show any impact on the inhibition of NANOG by GMO PMO antisense oligo treatment; however, treatment with chlorpromazine inhibited the morphological alteration and suppression of NANOG by NANOG **ASO GMO PMO**, indicating the involvement of clathrin mediated endocytosis pathway in the cellular uptake of antisense oligo (**Figure 6 G, S 5 C** and **S 6**). It is worthy to mention that dose of chloroquine, amiloride, genistein and chlorpromazine was applied to establish the mechanism didn’t show any morphological changes in MCF7 cells (**Figure S 7 A**). Moreover, MTT data didn’t show any cytotoxicity (**Figure S 7 B** and **Figure S 8 A**) and inhibition of NANOG (**Figure S 8 B**) at the respective concentrations.

### Inhibition of NANOG expression decreases stemness in PC3 and MDA MB-231 cells

The efficacy of NANOG **ASO GMO-PMO 6** was further evaluated in prostate cancer cell line PC3 and triple negative breast cancer cells MDA MB-231 also. The immunoblot data showed the dose dependent decrease of NANOG expression in PC3 cells. The effect was more prominent with 2 μM dose of NANOG **ASO GMO PMO 6** and therefore used in rest of the experiments (**Figure S 9 A, B**). The MTT data showed significant decrease in cell viability at a dose of 70 μM of scrambled oligo GMO PMO while the same result was obtained with 1 μM dose of NANOG ASO GMO PMO (**Figure S 9 C, D**), indicating negligible toxicity of GMO PMO chimera at lower doses for PC3 cells also. The expression of several stemness factors like NANOG, Sox-2, c-Myc and Cxcr4 were detected by immunoblot. NANOG **ASO GMO-PMO** suppressed the expression of these stemness related genes (**Figure 7 A** and **Figure S 10 A**) and also other EMT responsible genes such as N-Cadherin, Vimentin, MMP-9, Snai1 and Twist (**Figure 7 A** and **Figure S 10 B, C**) while increased the E-Cadherin level. The suppression of NANOG also increased apoptosis in PC3 cells as reflected by higher Bax expression with lower Bcl2 level (**Figure 7 A**, **Figure S 10 D**). The results were further confirmed by migration wound assay as well as clonogenicity assay where higher effect was seen in case of NANOG **ASO GMO PMO 6** compared to taxol and NANOG ASO treatment (**Figure S 11 A-D**).

**Figure 7:**
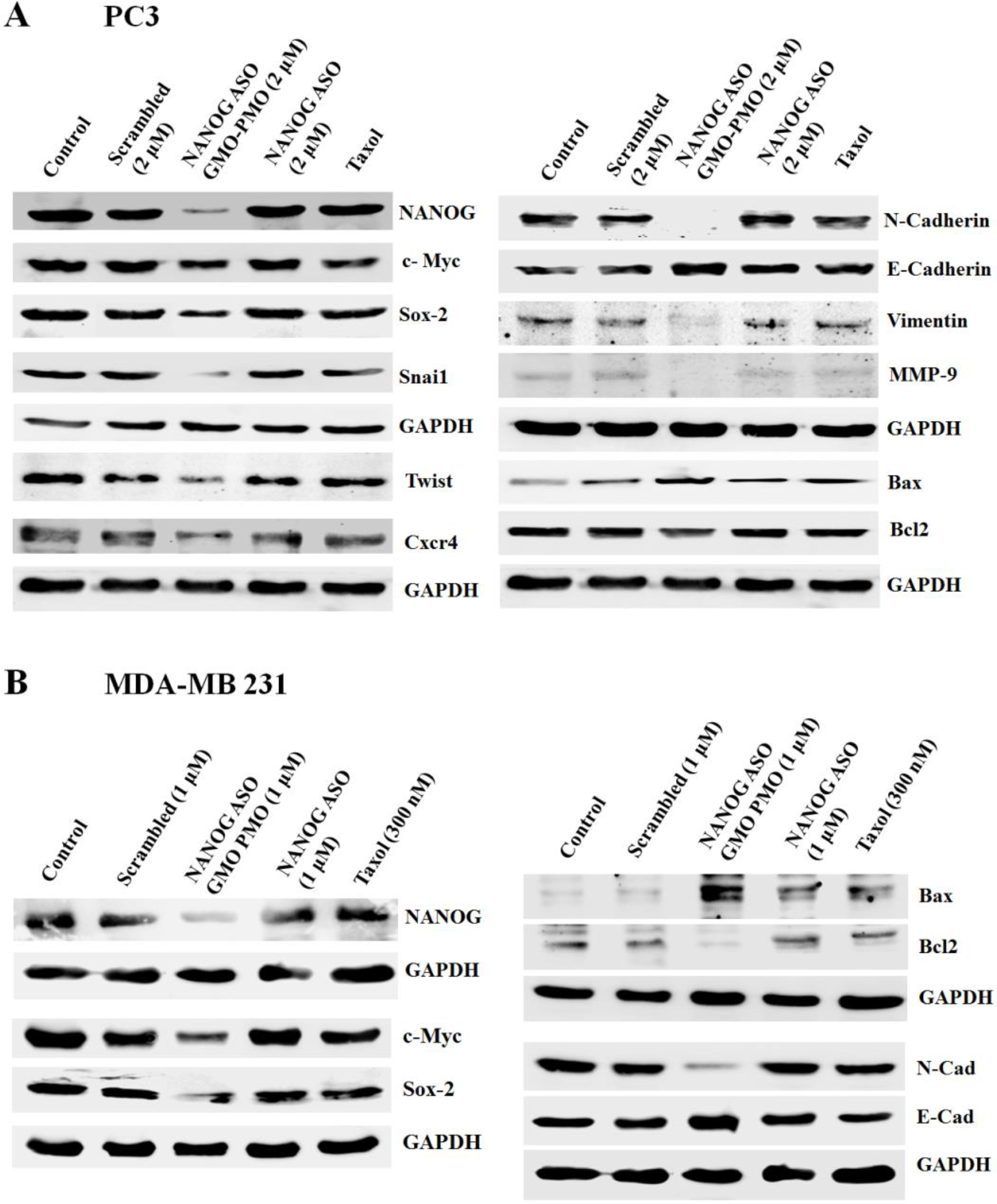
A. Immunoblot data show the expression of NANOG, Sox-2, Cxcr4, c-Myc, EMT markers such as N-Cadherin, Vimentin, Snai1, Twist, MMP9, anti-apoptotic Bcl2 and pro apoptotic protein Bax in PC3 cells. B. Immunoblot data show the expression of NANOG, Sox-2, c-Myc, EMT markers such as N-Cadherin, E-cadherin, anti-apoptotic Bcl2 and pro apoptotic protein Bax in MDA MB-231 cells. The protein expression levels are expressed relative to GAPDH. NANOG ASO stands for naked PMO from Gene Tools.

The MDA MB-231 cells were treated with increasing doses of NANOG **ASO GMO-PMO** and scrambled NANOG GMO-PMO. The MTT assay showed significant decrease in cell viability at a dose of 90 μM of scrambled GMO-PMO whereas the viability was significantly reduced with 0.5 μM dose of NANOG **ASO GMO-PMO**(**Figure S 12 A, B**), showing very less toxicity of chimera at lower doses for triple negative breast cancer cells also. The immunoblot data showed a dose dependent decrease of NANOG expression and 0.5, 1 and 2 μM doses of chimera showed similar level of NANOG inhibition in MDA MB-231 cells (**Figure S 12 C, D**) and 1 μM dose of NANOG **ASO GMO PMO 6** was used in rest of the experiments. The inhibition of NANOG (**Figure 7 B** and **Figure S 12 E**) also suppressed the other stemness factors c-Myc, Sox-2, EMT markers N-cadherin while increased E-cadherin level (**Figure 7 B** and **Figure S 12 E**). Suppression of NANOG increased the apoptosis of MDA MB-231 cells as depicted by higher Bax expression with lower level of Bcl2 anti-apoptotic proteins (**Figure 7 B** and **Figure S 12 E**).

### NANOG ASO GMO PMO 6 induced NANOG knockdown enhances chemo-sensitivity to taxol

One of the major challenges in cancer treatment that is accountable for poor treatment outcomes and tumor relapse is the emergence of multidrug resistance. The molecular mechanisms involved in such chemoresistance are the over expression of ATP Binding Cassette (ABC) transporters mainly that of mdr1 and ABCG2 which is specifically over expressed in Cancer Stem Cells [44‒46]. Since, NANOG has been reported to confer chemoresistance in cancer cells, this study was done to elucidate whether NANOG inhibition can enhance drug sensitivity of cells to taxol.

The cells were treated with **scrambled Nanog 7**, 0.5 μM and 1 μM of NANOG ASO GMO PMO and NANOG ASO for 36 h and each of them were subjected to 12.5 nM, 25 nM, 50 nM and 100 nM doses of taxol for 48 h and 96 h, respectively. The cell viability by MTT assay indicated that NANOG knocked down cells exhibited increased sensitivity to taxol thus successfully increasing drug sensitivity to taxol in MCF7 cells (**Figure 8 A, S 13 A, B**). To find out the reason, we evaluated the expression of drug resistance genes MDR1 and ABCG2 by immunoblot. The results showed significant down-regulation of MDR1 and ABCG2 in NANOG silenced cells (**Figure 8 B, S 14**). The results were further confirmed by immunocytochemistry where significant reduction in the expression of MDR1 was obtained (**Figure 8 C** and **S 15, 16**). Since higher rate of inhibition was obtained in case of combined treatment of NANOG **ASO GMO PMO** and Taxol, the rest of the experiments were executed with the combination treatment.

**Figure 8:**
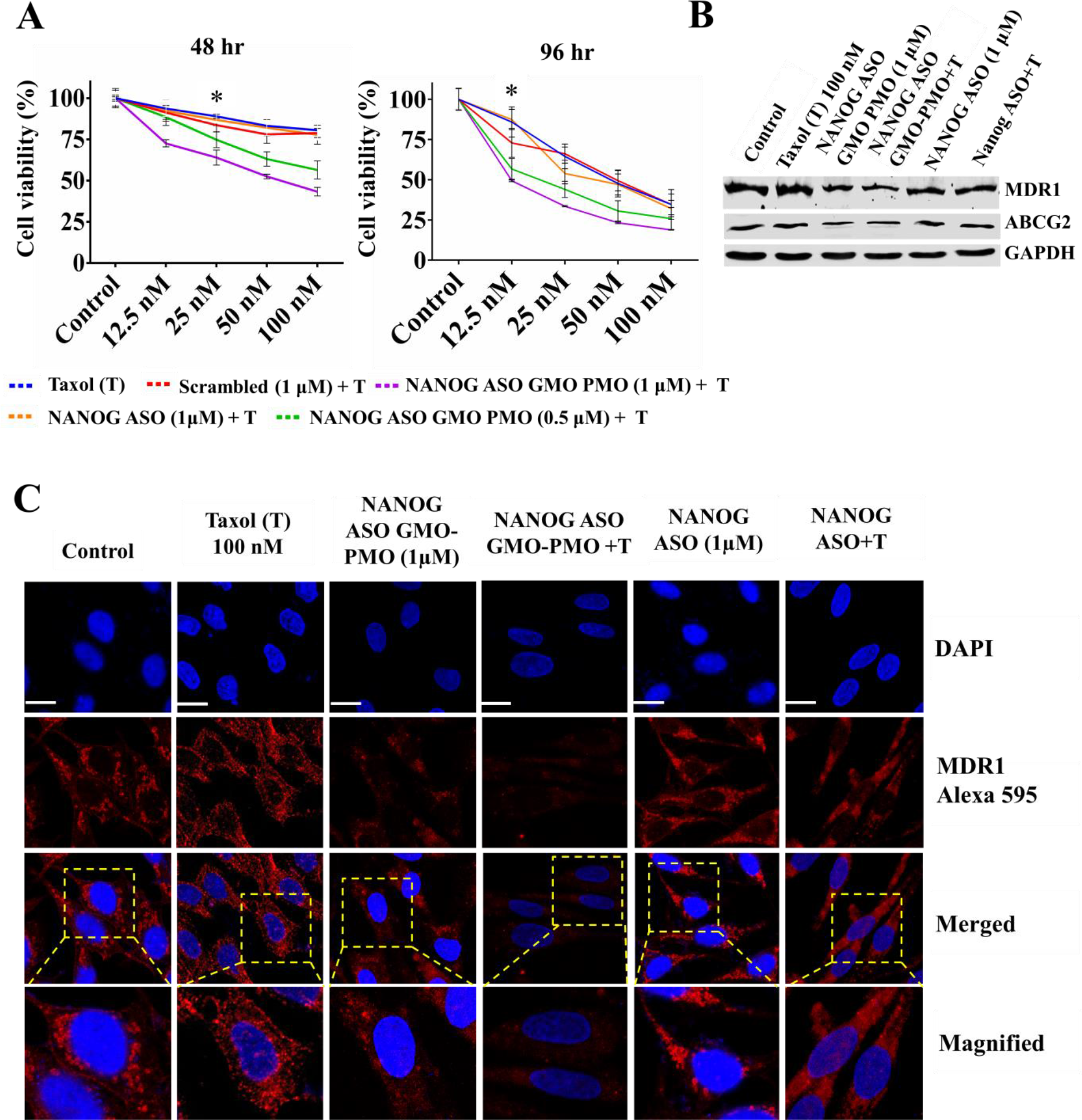
A. Different antisense treated MCF7 cells were subjected to 12.5nM, 25nM, 50nM and 100nM doses of Taxol and cell viability was measured by MTT assay after 48 and 96 h respectively. B. NANOG knocked down cells also showed lower expression levels of MDR1 and ABCG2 as depicted from immunoblotting. C. Confocal immunofluorescence microscopic analysis of Mdr1 (shown in red) in control and treated cells after 72 h of combined treatment with antisense and Taxol. Nuclei were stained with hoechst (blue). Quantification of MDR1 intensity per nucleus obtained from confocal immunofluorescence microscopy was calculated for 20–25 cells. NANOG ASO stands for naked PMO.

### Taxol promulgates apoptotic response in NANOG silenced cells

The expression of Bax, a pro-apoptotic signalling molecule, was examined by immunoblotting. It was found significantly higher in NANOG silenced cells and effect was more prominent in combination group. In contrast, both the NANOG knocked down cells and combination group showed lower Bcl2 expression which indicated higher level of Bax/Bcl2 ratio to sustain pore formation in mitochondria and altered the membrane potential (**Figure 9 A** and **S 17**). The leakage of CytC due to the loss of membrane potential aggravated the activation of intrinsic Caspase pathway which was reflected by the higher activation Caspase 9 in combination group (**Figure 9 B**). Moreover, the TUNEL formation again confirmed the activation of DNA endonuclease which ultimately cleaved the DNA. The numbers of TUNEL positive cells were higher in combination group compared to NANOG silenced cells and taxol treated cells which indicated that silencing of NANOG by NANOG **ASO GMO-PMO** sensitizes the MCF7 cells towards taxol and thus, increased the efficacy of taxol by modulating the chemosensitivity of MCF7 cells (**Figure 9 C** and **S18-20**).

**Figure 9:**
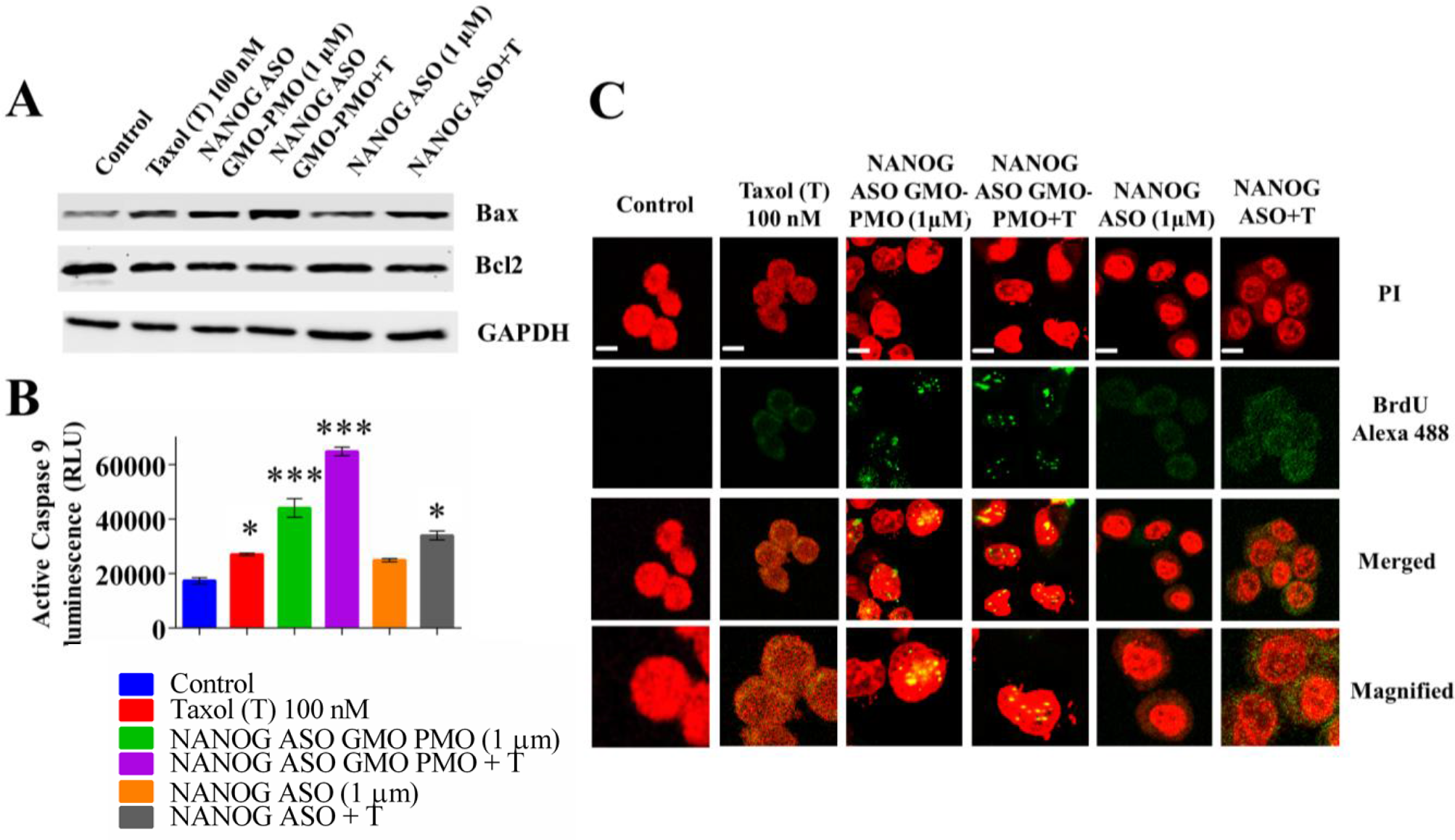
A. Taxol increases apoptosis of the NANOG suppressed MCF-7 cells. The immunoblot data show the Bax and Bcl2 level in MCF-7 cells. B. The bar diagram shows the active Caspase 9 level. C. The BrdU incorporation assay to detect the apoptosis in MCF-7 cells. The Nuclei were stained by PI and the incorporation of BrdU was detected by the alexa 488 tagged (green) anti BrdU antibody. Quantification of TUNEL formation/nucleus was obtained from confocal immunofluorescence microscopy and was calculated for 20–25 cells. Error bars indicate means ± SE (n = 3), and data are presented as percentages relative to the non-treated MCF-7 cells. *p< 0.05.

### GMO PMO Mediated Nanog Regulation in MCF10A

MCF10A is an immortalized non transformed epithelial mammary cell line which has been used as a control over here. The way MCF7 was subjected to treatment with different doses of scrambled GMO PMO, likewise MCF10A too was treated with the same to ensure whether there is any effect. From the cell viability assay, it was found that scrambled GMO PMO had no effect on MCF10A upto100 μM dose (**Figure 10 A**). Not only that, MCF10A also did not show any effect in MTT assay when treated with 0.5, 1 and 2 μM of NANOG **ASO GMO PMO**(**Figure 10 B**). The same result was obtained from immunoblotting where higher expression of NANOG was seen in MCF7 and PC3 and their corresponding inhibition w.r.t. MCF10A (**Figure 10 C, D**). Inspite of getting negligible inhibition of NANOG in MCF10A, we were intrigued to see whether NANOG ASO GMO PMO had any effect on other reprogramming factors. It is evident from the western blot data that the antisense oligo treatment did not alter the expression of NANOG, Sox-2, Oct4 and c-Myc stemness factors in MCF-10A thereby confirming its specificity in case of cancer cells (**Figure 10 E, F**).

**Figure 10:**
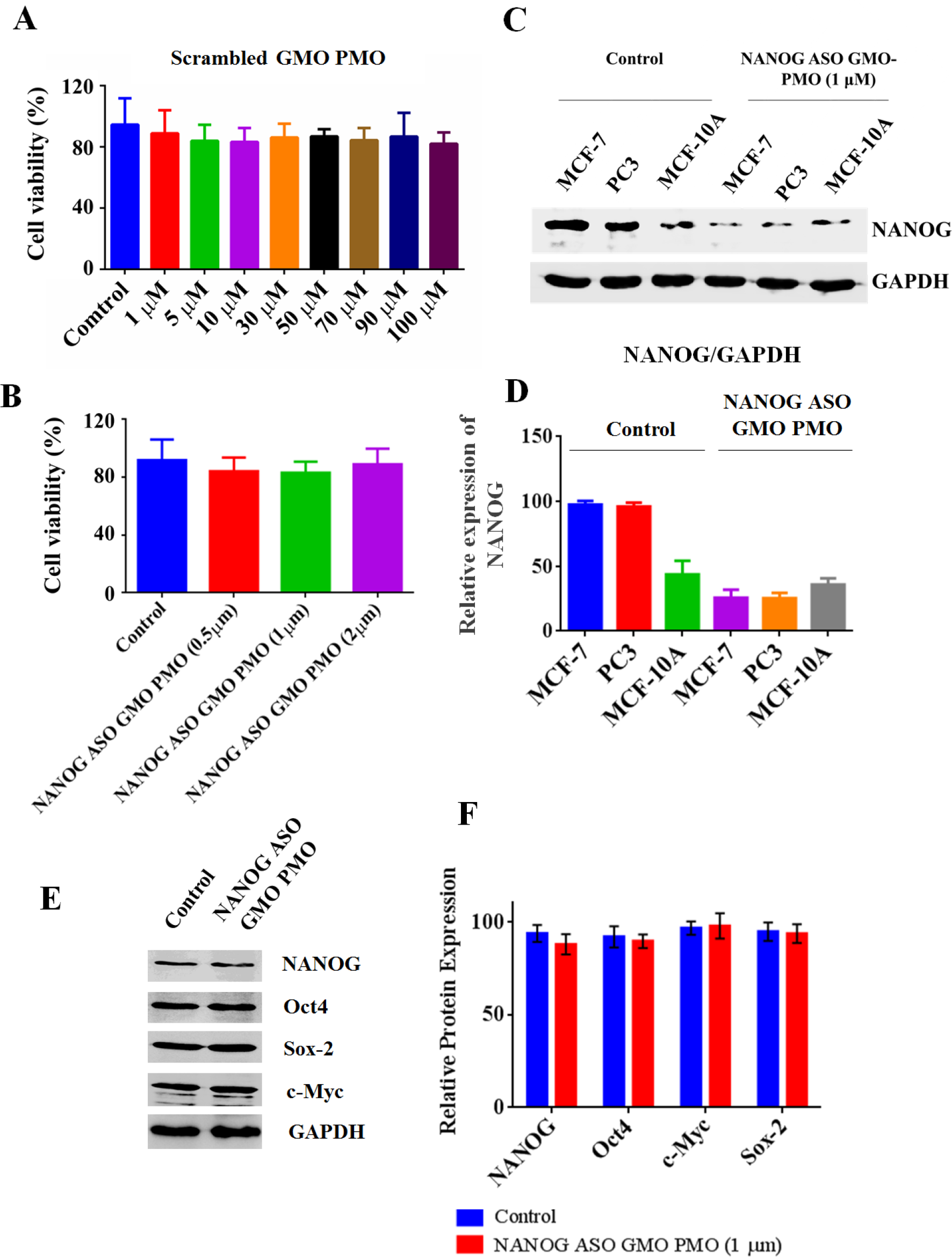
The NANOG ASO GMO-PMO does not has any effect on MCF-10A. (A) Cell viability assay (MTT assay) of MCF10A after treatment with different doses of scrambled GMO PMO and (B) NANOG ASO GMO PMO. (C) Western blot analysis showing the expression of NANOG in MCF7, PC3 and MCF10A before and after treatment with NANOG ASO GMO PMO. (D) The densitometric analysis of immunoblot data. (E) The immunoblot data of NANOG, Oct4, Sox-2 and c-Myc in MCF-10A. (F) The densitometric analysis of the immunoblot data. The protein expression levels are expressed relative to GAPDH. Error bars indicate means ± SE (n = 3), and data are presented as percentages relative to the non-treated MCF-7 cells. *p< 0.05.

## Conclusion

Antisense oligonucleotides (ASO) are rapidly emerging as a therapy of choice for genetic disorder diseases [47–49]. As of January 2020, ten oligonucleotides-based drugs have been approved by the FDA in which Eteplirsen [4] and Golodirsen [5] are based on PMO. However, in order to improve the therapeutic efficacy, the delivery vehicle is essential for the delivery of PMO and for other ASO in general [7, 47–49] and accordingly guanidinium rich cellular transporters conjugated PMOs have been developed [8]. In this context, our self-transfecting GMO-PMO or PMO-GMO chimera concept is new and novel where there is no need of using any delivery vehicle. Unlike fluorescent-labelled VivoPMO/PPMO/IGT-PMO synthesis, its Bodipy conjugation synthesis has been achieved easily for localization study in *in vivo* zebrafish model. Though PMO mediated inhibition of specific genes in malignancy had already been reported in *in vitro* and *in vivo* studies [50–52], however, PMO-therapy has not been approved for cancer treatment though Phase I study was conducted successfully [50]. In our case the delivery of PMO is much easier than previously reported method which perhaps was the rate limiting step for bringing PMO therapy for cancer treatment.

Since NANOG plays a crucial role in the stemness, growth of proliferation and chemoresistance of various types of malignancy such as breast, ovarian, bladder, gastric, melanoma and many others, we have targeted to knock down NANOG with our cell penetrating GMO-PMO chimera. Additionally, most of the chemotherapeutic drugs have limited efficacy and act non-specifically which led to the development of drug resistance in tumour cells. The silencing of NANOG leads to the ablation of CSC self-renewal, cell proliferation and growth. Interestingly, when taxol was added to the NANOG silenced cells, it led to the aggravation of the effect of taxol which was higher than Taxol treatment alone. The probable explanation of getting this effect is that the silencing of NANOG led to the decrease in the expression of Mdr1 and ABCG2 genes thereby, increasing the chemosensitivity of MCF7 cells towards taxol. Moreover, silencing of NANOG increases the Bax/Bcl2 ratio which sustains the leakage of Cyt C from mitochondria and activates apoptotic pathway as observed by higher level of active Caspase 9 and TUNEL formation in combination over the taxol treatment alone. Therefore, the antisense technology has emerged as a promising strategy for disease relevant gene silencing on a rational basis and accordingly, we evaluated the efficacy of NANOG **ASO GMO PMO** as a therapeutic agent against stemness factor NANOG which affected the associated oncogenic pathways to prevent the cancer progression.

The Ntl GMO PMO showed remarkable inhibition of tail formation in zebrafish embryo which indicates the successful targeted *in vivo* delivery of antisense PMO. It is worth mentioning that GMO-PMO works even after 8-cell stage which was not possible to achieve earlier using regular PMO except using photocaged PMO [53–57] or our IGT-PMO [**15**], perhaps could be useful for the regeneration biology in zebrafish model. To examine the species specific efficacy of GMO PMO chimera, Gli1 ASO GMO PMO was synthesised to target the murine Gli1 protein in ShhL2 cells and the treatment with Gli1 ASO GMO PMO substantially decreased the Gli1 expression and other related stemness factors in ShhL2 cells. The same protein has also been targeted earlier using IGT conjugated PMO (IGT-PMO) [16] was found to down regulate Gli protein at 5 μM dose. As GMO-PMO showed relatively higher down-regulation of Gli1 with a lower dose (750 nM), hence, GMO-PMO is 6.6 times more effective than IGT-PMO in gene silencing activity. It is important to mention here that among the cell-permeable PMOs, Pip6a-PMO was the most efficient conjugate to date which worked in nanomolar dose where Pip6a represents the modified CPP having the sequence Ac-RXRRBRRXRYQFLIRXRBRXRB-OH, with B = β-alanine and X=amino hexanoic acid [58]. In the case of GMO-PMO/PMO-GMO chimeras, no such delivery vehicle is required and showed the antisense efficacy as low as 750 nM dose in *in vitro* and required as minimum as four guanidinium linkages. Though, backbone modified PMO and their chimeras [59, 60] or PMO-DNA chimera [61] or chimeric oligonucleotides containing morpholino thymidine analogues [62, 63] or with a single guanidinium morpholino unit incorporated DNA [64] has been reported earlier however, to the best of our knowledge, GMO-PMO/PMO-GMO chimera is the first report from our group. Considering the simplicity, the possibility of rational design, relatively inexpensive and easy synthesis, GMO-PMO/PMO-GMO self-transfecting chimera, in principle could be useful as tools to target any gene including virus and bacteria towards the development of antisense therapy and could be a practical approach. Their high level of antisense efficacy in a low dose (500-750 nM) could overcome the longstanding problems associated with off target effects of antisense reagents [49]. The present study has intended to use ASO GMO PMO chimera as a targeted gene silencing agent which could be a potential therapeutic regime in near future.

## Experimental

### Materials and Methods

The MCF-7, PC3, MCF10A and MDA MB-231 were purchased from NCCS, Pune, India. The antibodies used in immunoblot and immunofluorescence study were procured from Cell Signaling Technology (CST) and Sigma Aldrich. The DMEM, FBS and antibiotic cocktail were taken from Gibco. The Caspase-Glo 9 Assay kit was purchased from Promega. The APO-BrdU TUNEL Assay Kit were obtained from Thermo Fisher Scientific. All the oligos used were > 95% pure in HPLC.

#### Microinjections into zebrafish embryos and imaging

Embryos used in these studies were obtained by natural mating of adult zebrafish (wild type) and staged. The embryos were cultured in E3 embryo medium at 28.5°C according to standard procedures. Microinjection of zebrafish was done using Eppendorf femtojet express microinjection setup. Compounds were microinjected in the embryos at the cell-yolk interface in different stages. 9 ng (0.25 mM stock, 3‒4 nl) of morpholino was injected into zebrafish embryo at 16 cell stage (1.6 hpf), 32 cell stage (1.75 hpf) and 64 cell stage (30‒33 embryos were injected in each set). H2O injected embryos were considered as control. Injected embryos were then transferred to Petri dishes and cultured at 28.5°C. Fluorescence intensities and distribution in the larvae were observed under an inverted fluorescence microscope (Olympus IX51) [16]

#### Cell Culture and treatment

Shh-Light 2 (derived from mouse embryo NIH3T3 cell lines stably transfected with a Gli-dependent firefly luciferase and constitutive renilla luciferase reporters) cells and HEK 293T cells overexpressing Shh-N (N-terminal fragment of Shh without cholesterol modification) cells were obtained as a gift from Professor J. K. Chen, Stanford University. The culture of Shh-Light 2, MCF7, PC-3, MDA MB-231 cells was performed in 10 % bovine calf serum (Gibco)–DMEM supplemented with 100 μg/mL streptomycin and 100 units/mL penicillin at 37° C in a 5% CO2 incubator in humidified atmosphere, trypsinized with 0.25 % trypsin– EDTA. The Shh-N cells were cultured to 80 % confluency in DMEM supplemented with 10 % FBS, 100 μg/ml S-25 streptomycin and 100 units/mL penicillin. The media was replaced with 2 % FBS–DMEM containing 100 μg/ml streptomycin and 100 units/ml penicillin and cultured for another 24 hr. and the media was collected, centrifuged and filtered through 0.22 μm filter. The Shh-N conditioned media was stored at –20°C and used to activate Shh pathway in Shh-Light 2 cells [**20**].

After the seeded cells reached ~ 50% confluency in separate culture plates, the Gli1/NANOG-specific ASO GMO PMO chimera, multi-mutated GMO PMO (MM)/Scrambled (NANOG) GMO-PMO, Gli1 ASO (PMO)/ NANOG ASO and known Hh inhibitor Cyclopamine/ chemotherapeutic drug taxol were added and incubated for 36 hours [20].

#### Cell viability assay

The viability of the MCF7, PC-3, MDA MB-231 and MCF10A was studied with scrambled GMO-PMO and NANOG ASO GMO-PMO to observe the toxicity of the chimera and the effect of ASO GMO-PMO, respectively. The viability of Shh-L2 cells with MM GMO PMO was studied to observe the toxicity of the chimera. The cells were seeded in 96-well plates at a density of 1× 10^4^ cells/well for 24 h before treatment with different concentrations of oligonuleotides. Post 72 h treatment, 100 μl of MTT solution was added to each well. After 4 hrs the media containing MTT was removed and the purple formazan product was dissolved in 100 μl/ well of DMSO and quantified by plate reader at 570 nm [15]

#### Western Blotting Analysis

Protein extracts from compound treated cells were resolved by 8-10% SDS polyacrylamide gel electrophoresis, transferred to nitrocellulose membranes and probed with primary antibodies against the proteins. The membranes were then subsequently probed with a horse radish peroxidase-conjugated secondary antibody and developed with Femto in Biorad Chemidoc [15].

#### Immunocytochemistry

Cells were grown on coverslips (pre-treated with Poly L Lysine), fixed with 4% paraformaldehyde for 10mins, permeabilized with 0.5% Triton X-100 for 5 min, then blocked in 1% BSA in PBS for 1 hr and incubated with primary antibodies overnight at 40C. Cells were then stained with Alexa 488 anti-rabbit IgG or Alexa 594 anti-mouse for 1 hr at RT, washed twice in PBS, nuclei stained with DAPI which were then mounted on Slow Fade Gold antifade reagent and visualized under Carl Zeiss confocal microsope (Holdtecs Technology Co., Ltd, Chengdu, China) [15].

To check the subcellular localization of NANOG **ASO GMO-PMO**, bodipy tagged GMO-PMO antisense oligo (0.5 μM) was added to MCF7 cells and incubated for 5 h and 18 h. The cells were counter stained with Mitotracker red and Lysotracker red separately for 1 hr and observed under Carl Zeiss confocal microsope after fixation in 4% paraformaldehyde [15].

#### Flowcytometry

To understand the cell transfection efficiency of NANOG **ASO GMO-PMO**, the MCF7 cells were treated with 0.5 and 1 μM doses of bodipy tagged NANOG **ASO GMO-PMO** and incubated for different time points. The cells were also treated with different doses of bodipy tagged NANOG **ASO GMO-PMO**(50-500 nM) and incubated for 4 h. The fluorescence intensity of MCF7 cells were measured by flow cytometry (BD FACS ARIA III) against untreated control. The fluorescence of 1×10^4^ cells in individual group was recorded [15].

#### Colony Formation Assay

Briefly, cells were grown in 6 well plates, treated with compounds and kept at 37°C in incubator for 15 days with intermittent media changes. After the stipulated time, cells were then stained with 0.005% crystal violet for 1 h and images were taken under a microscope [15].

#### Wound Healing Assay

The MCF7 and PC-3 cells were cultured to completely confluent state in 12-well plates and a scratch was made across the cell monolayer of each sample well with a 10 μl pipette tip. Then the cell monolayer was washed with PBS and incubated with compounds in 0.5% FBS DMEM at 37°C with 5% CO2 for 36 h. The widths of the scratches were measured and the percentages of relative wound closure were compared at 0h and 36h, respectively [15].

#### Chemo-sensitivity Assay

The MCF-7 cells were seeded in 96 well plates at a density of 1.0×10^4^cells/well for 24 h. Before taxol addition at concentrations of 12.5, 25, 50 and 100 nM, cells were separately treated with NANOG ASO GMO-PMO, scrambled GMO-PMO and NANOG ASO for 36 h. After addition of different concentrations of taxol, the cell viability was evaluated by MTT assay at different time points (48h and 96h respectively) [**15**].

#### Caspase 9-Glo assay and TUNEL assay

Combination treatment was conducted to check the effect of NANOG treatment on the chemosensitivity of MCF7 cells towards Taxol. The cells were treated with PMO (NANOG ASO), and NANOG ASO GMO PMO for 36 h and then they were again subjected to taxol treatment (100 nM) for another 36 hr. The combination treated cells were studied for the expression of several apoptotic proteins by fluorescence and immunoblot techniques. The activity of Caspase 9 was measured by Caspase-Glo 9 Assay Systems of promega kit and the TUNEL formation was measured according manufacturer protocol of APO-BrdU TUNEL Assay Kit of Thermofisher.

## Author Contributions

SS: Conceived the idea, designed the hypothesis. SS and UD wrote the manuscript, JK: Synthesized morpholino monomers, oligomers, purified and characterized. U.D and C.B: Biology work. J.B: Synthesized the no tail GMO-PMO and biological experiments in zebrafish embryos.

## Notes

The authors declare no competing financial interest.

## ACKNOWLEDGMENT

J.K., U.D. and C.B. thank IACS for their fellowship. All authors have given approval to the final version of the manuscript.

## ASSOCIATED CONTENT

**Supporting Information**. Detailed experimental procedures and characterization data including the spectra for all new compounds can be found in the supporting information.

