## Supplemental Data for "Self-transfecting GMO-PMO and PMO-GMO chimeras enable gene silencing *in vitro and in vivo* zebrafish model and NANOG Inhibition Induce the Apoptosis in Breast and Prostate Cancer Cells"

**Table of Contents** **Page**

General Information.......................................................................................................... S-2

HPLC chromatograms of **GMO-PMOs 1 to 7**..................................................................S-2-7

Supporting figure 1: Bodipy-no tail phenotypes of zebrafish image……………..................... S-07

Supporting figure 2: Cytotoxicity and dose standardization of GMO-PMO………………….S-08

Supporting figure 3: Serum and time dependent effect of NANOG ASO GMO-PMO…….....S-09

Supporting figure 4: Cell transfection and sub-cellular localization………………………….S-10

Supporting figure 5: Densitometric analysis…………………………………………………...S-11

Supporting figure 6: Morphological alteration of MCF-7 cells……………………………S-12-13

Supporting figure 7, 8: Morphological alteration, cytotoxicity with inhibitors alone….….S-14-15

Supporting figure 9: Cytotoxicity and dose standardization in PC3 cells…………….........S-16

Supporting figure 10: Densitometric analysis of PC3 immunoblot data….…………………...S-17

Supporting figure 11: Wound healing and colony assay in PC3 cells………………………S-18-19

Supporting figure 12: Cytotoxicity and dose standardization in MDA MB-231 cells……..S-20-21

Supporting figure 13: Chemo-sensitivity assay………………………………………….S-21-22

Supporting figure 14: Densitometric analysis of MDR1 and ABCG2 immunoblot data……..S-22

Supporting figure 15: Immunofluorescence images of MDR1……………………………...S-23

Supporting figure 16: Enlarged images of merged data………………………………………..S-24

Supporting figure 17: Densitometric analysis of Bax and Bcl2………………………………..S-25

Supporting figure 18-20: Immunofluorescence images of TUNEL assay…………………S-26-28

### General Information

Zebrafish embryos were obtained by natural mating of adult zebrafish (wild type). The embryos were cultured in E3 medium at 28.5 $^{o}C$ according to standard procedure. Microinjection of zebrafish was done using Eppendorf femtojet express microinjection setup. * represents the guanidinium linkages in GMO. NANOG ASO stands for naked PMO from Gene Tools.

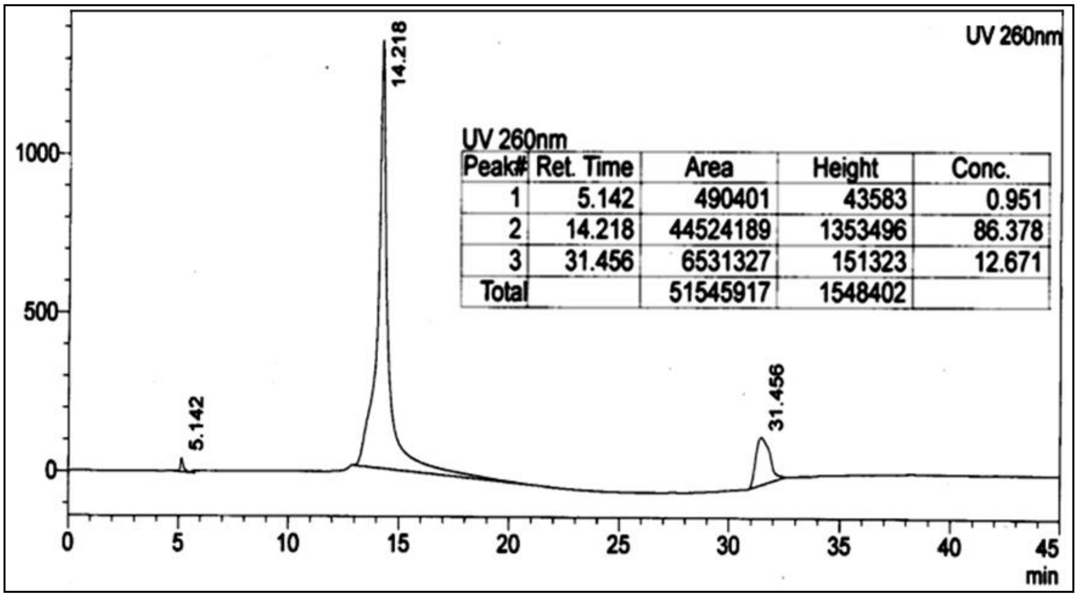


UV 260nm

| Peak# | Ret. Time | Area | Height | Conc. |
|---|---|---|---|---|
| 1 | 5.142 | 490401 | 43583 | 0.951 |
| 2 | 14.218 | 44524189 | 1353496 | 86.378 |
| 3 | 31.456 | 6531327 | 151323 | 12.671 |
| Total | | 51545917 | 1548402 | |

HPLC chromatogram of at-a-stretch Ntl **GMO-PMO-Ntl 1** chimera of sequence 5´-G*A*C*T*TG AGG CAG ACA TAT TTC CGA T-3´ (Gradient acetonitrile 0 to 35 % in 0.1 M $NH_4OAc$, 35 min, 1 mL/min).

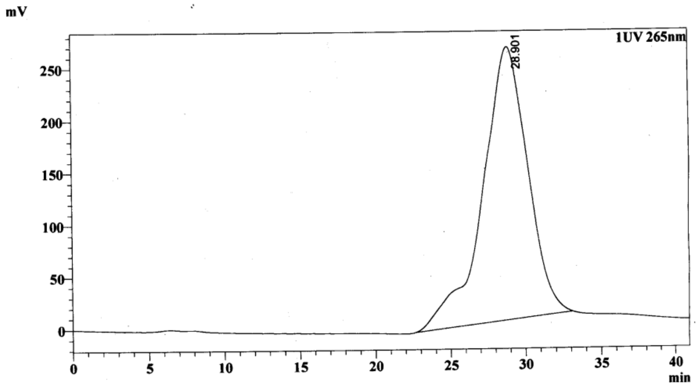


**Peak Table**

UV 265nm

| Peak# | Ret. Time | Area | Height | Area% |
|---|---|---|---|---|
| 1 | 28.901 | 54847018 | 262193 | 100.000 |
| Total | | 54847018 | 262193 | 100.000 |

HPLC chromatogram of at-a-stretch **GMO-PMO 1-BODIPY** chimera of sequence 5´- G*A*C*T*TG AGG CAG ACA TAT TTC CGA T-3´-Bodipy. (Gradient acetonitrile 0 to 35 % in 0.1 M $NH_4OAc$, 35 min, 1 mL/min)

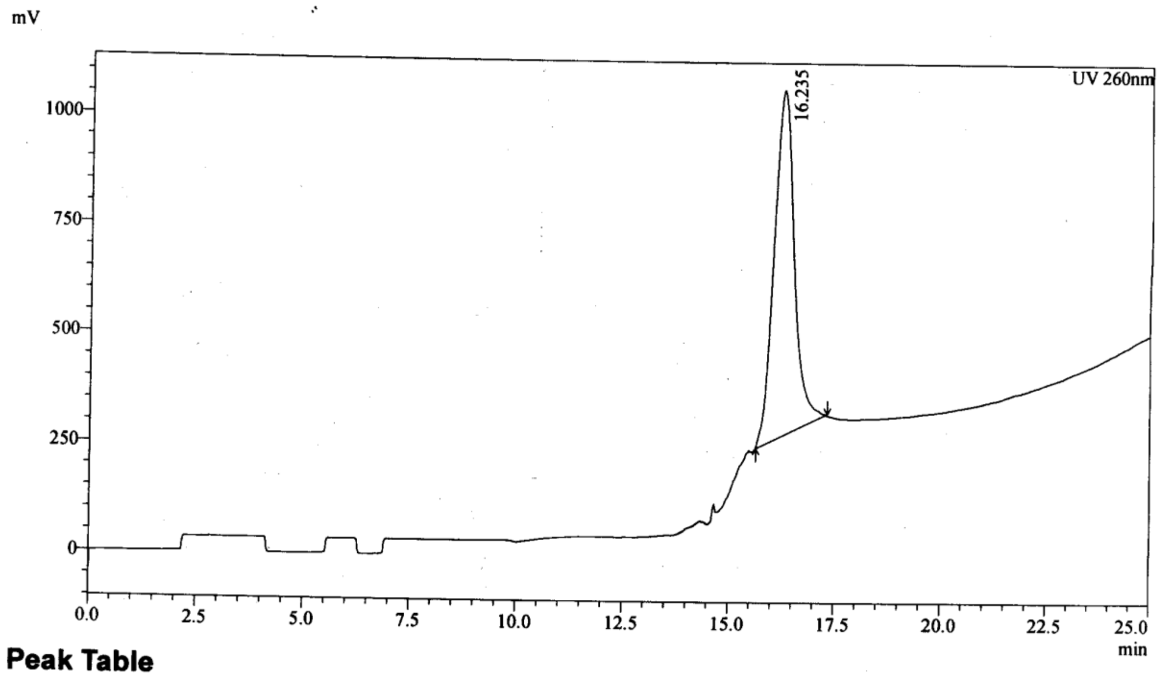


**Peak Table**

UV 260nm

| Peak# | Ret. Time | Area | Height | Area% |
|---|---|---|---|---|
| 1 | 16.235 | 26283298 | 792044 | 99.547 |
| 2 | 40.846 | 119660 | 55589 | 0.453 |
| Total | | 26402958 | 847633 | 100.000 |

HPLC chromatogram of at-a-stretch mGli **GMO-PMO 2** chimera of sequence 5´- T*T*G*GAT TGA ACA TGG CGT CT-3´ (Gradient acetonitrile 5 to 50 % in 0.1 M $NH_4OAc$, 25 min, 1 mL/min).

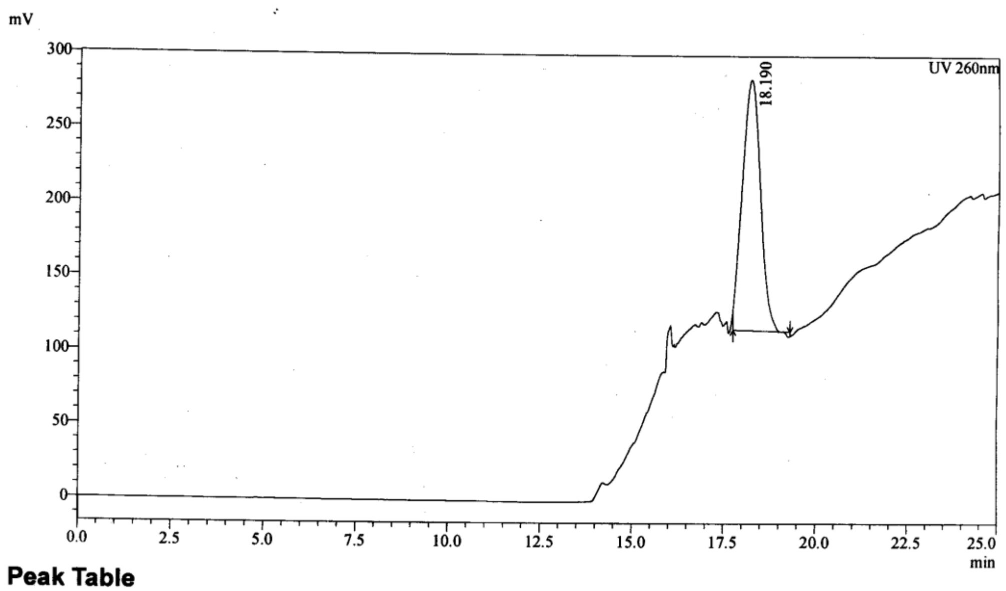


**Peak Table**

UV 260nm

| Peak# | Ret. Time | Area | Height | Area% |
|---|---|---|---|---|
| 1 | 18.190 | 5797123 | 169489 | 100.000 |
| Total | | 5797123 | 169489 | 100.000 |

HPLC chromatogram of at-a-stretch mGli single mismatched GMO-PMO **3** chimera of sequence 5′-T*T*G*T*AT TGA ACA TGG CGT CT-3′ (Gradient acetonitrile 5 to 50 % in 0.1 M $NH_4OAc$, 25 min, 1 mL/min).

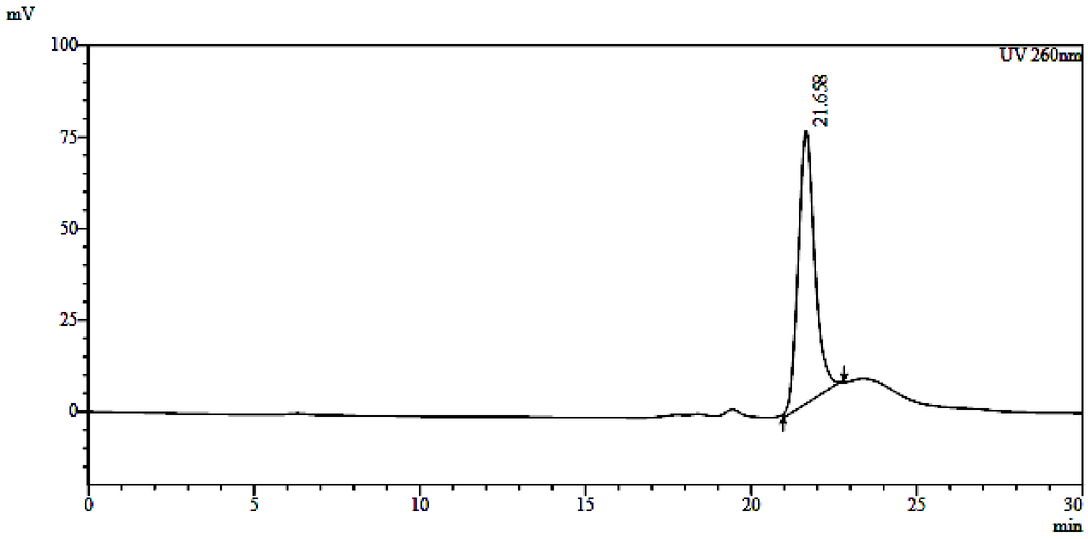


HPLC chromatogram of at-a-stretch mGli multi mismatched GMO-PMO **4** chimera of sequence 5′-G*G*T*T*AT TGA ACA TGG CGT CT-3′ (Gradient acetonitrile 5 to 50 % in 0.1 M $NH_4OAc$, 25 min, 1 mL/min).

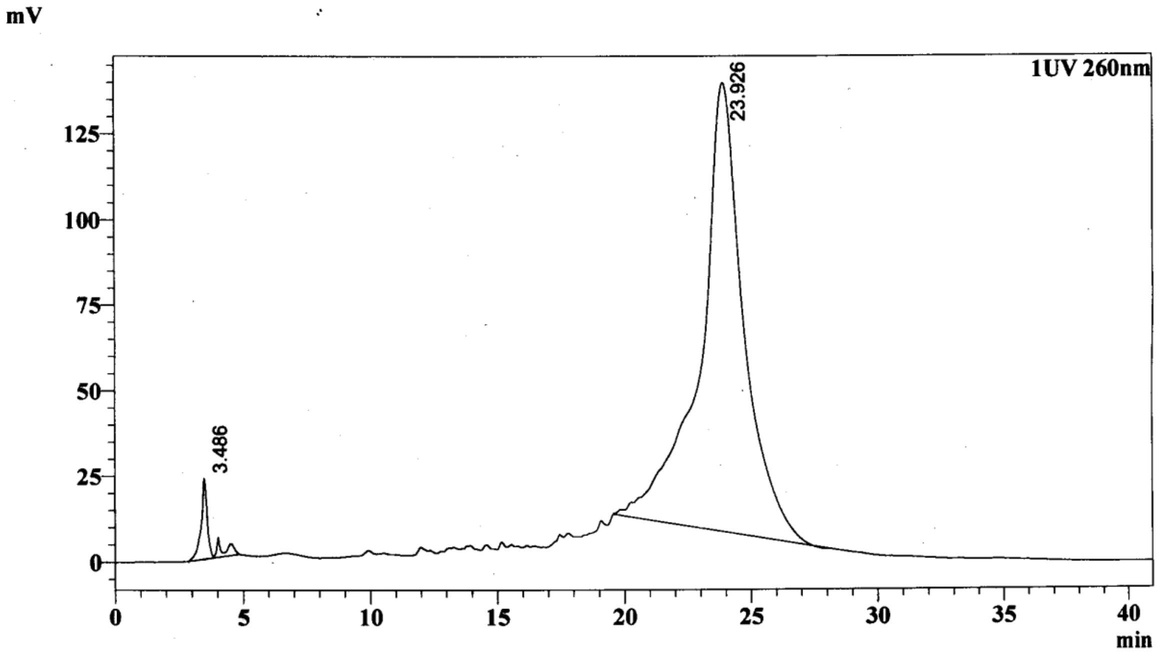


**Peak Table**

UV 260nm

| Peak# | Ret. Time | Area | Height | Area% |
|---|---|---|---|---|
| 1 | 3.486 | 510023 | 23396 | 3.117 |
| 2 | 23.926 | 15853595 | 131183 | 96.883 |
| Total | | 16363618 | 154579 | 100.000 |

HPLC chromatogram of at-a-stretch **mGli-PMO-GMO 5** chimera of sequence 5´- TTG GAT TGA ACA TGG C*G*T*C*T-3´ (Gradient acetonitrile 0 to 35 % in 0.1 M $NH_4OAc$, 35 min, 1 mL/min).

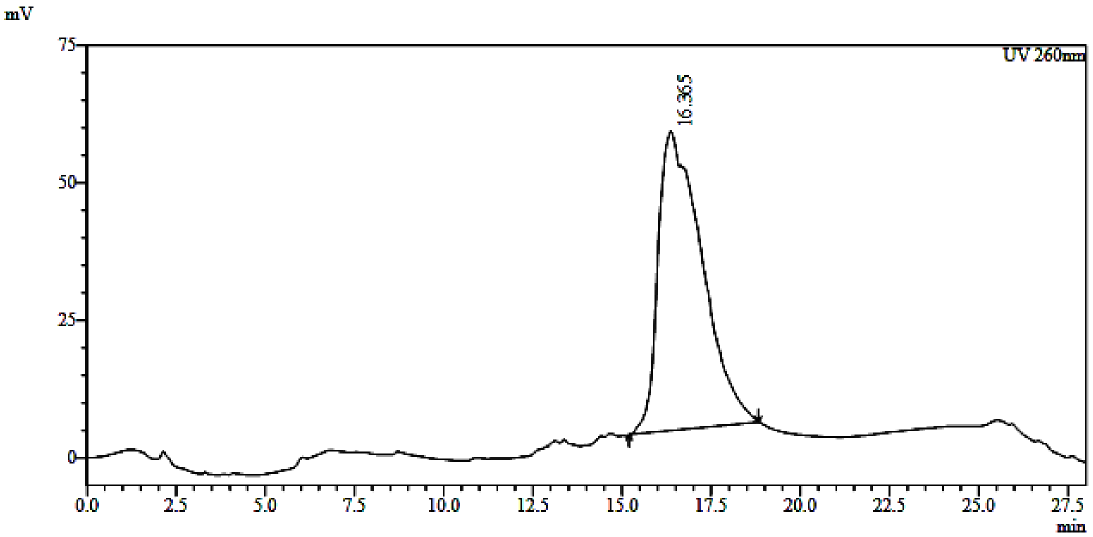


HPLC chromatogram of at-a-stretch Nanog **GMO-PMO 6** chimera of sequence 5′OH-G*T*G*A*GTTGCCTGCATAATAACATGA-3′ (Gradient acetonitrile 5 to 50 % in 0.1 M $NH_4OAc$, 30 min, 2 mL/min).

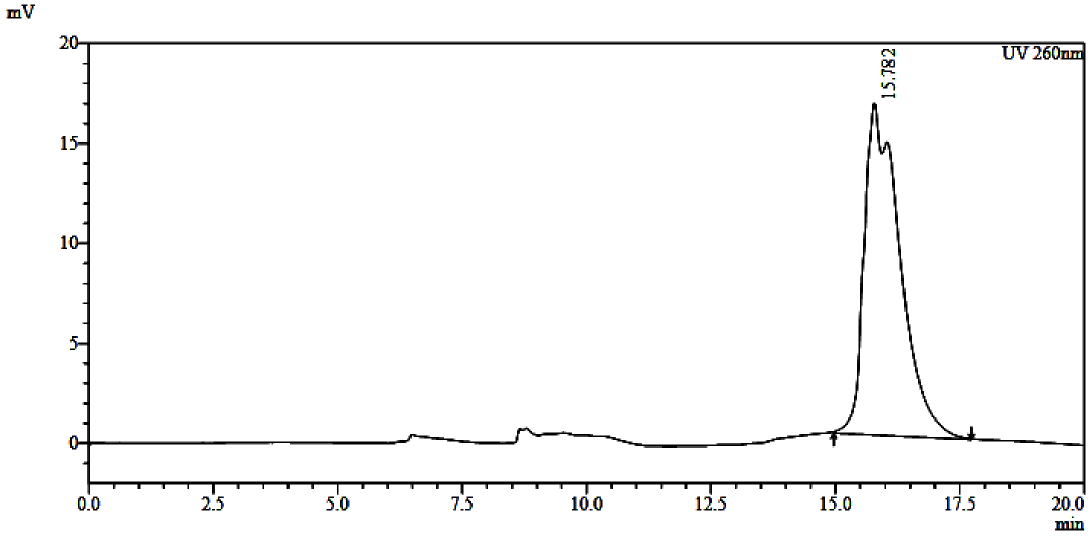


HPLC chromatogram of at-a-stretch scrambled-Nanog **GMO-PMO 7** chimera of sequence 5′OH-T*G*A*G*GTTCGCTGCATAATAACATGA-3′ (Gradient acetonitrile 5 to 50 % in 0.1 M $NH_4OAc$, 22 min, 2 mL/min).

**Biology results:**

**A**

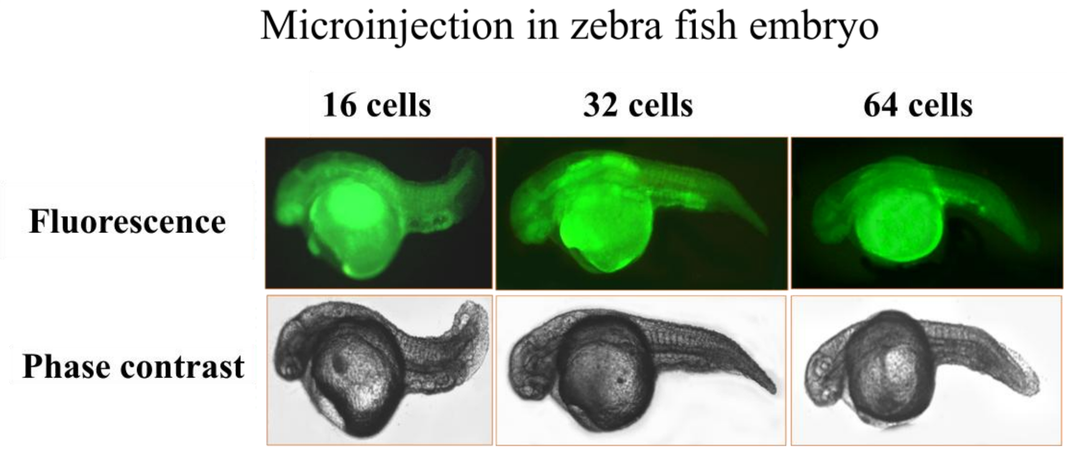


**B**

| Cells stages | Number of embryos injected | No of injected embryos alive | No of embryos showing phenotype |
|---|---|---|---|
| 2-4 | 30 | 27 | 25 |
| 4-8 | 31 | 26 | 24 |
| 8-16 | 33 | 28 | 25 |
| 16-32 | 32 | 26 | 23 |
| 32-64 | 30 | 25 | 21 |

**Supporting Figure 1: Zebrafish Bodipy-Ntl images. A.** The fluorescence images showed the distribution of Bodipy tagged *Ntl* antisense in zebrafish embryos. The phase contrast images of showed the phenotype of the respective embryos. **B.** Table showed the number of alive, phenotypically positive embryos among the Bodipy-Ntl injected embryos at different cells stages.

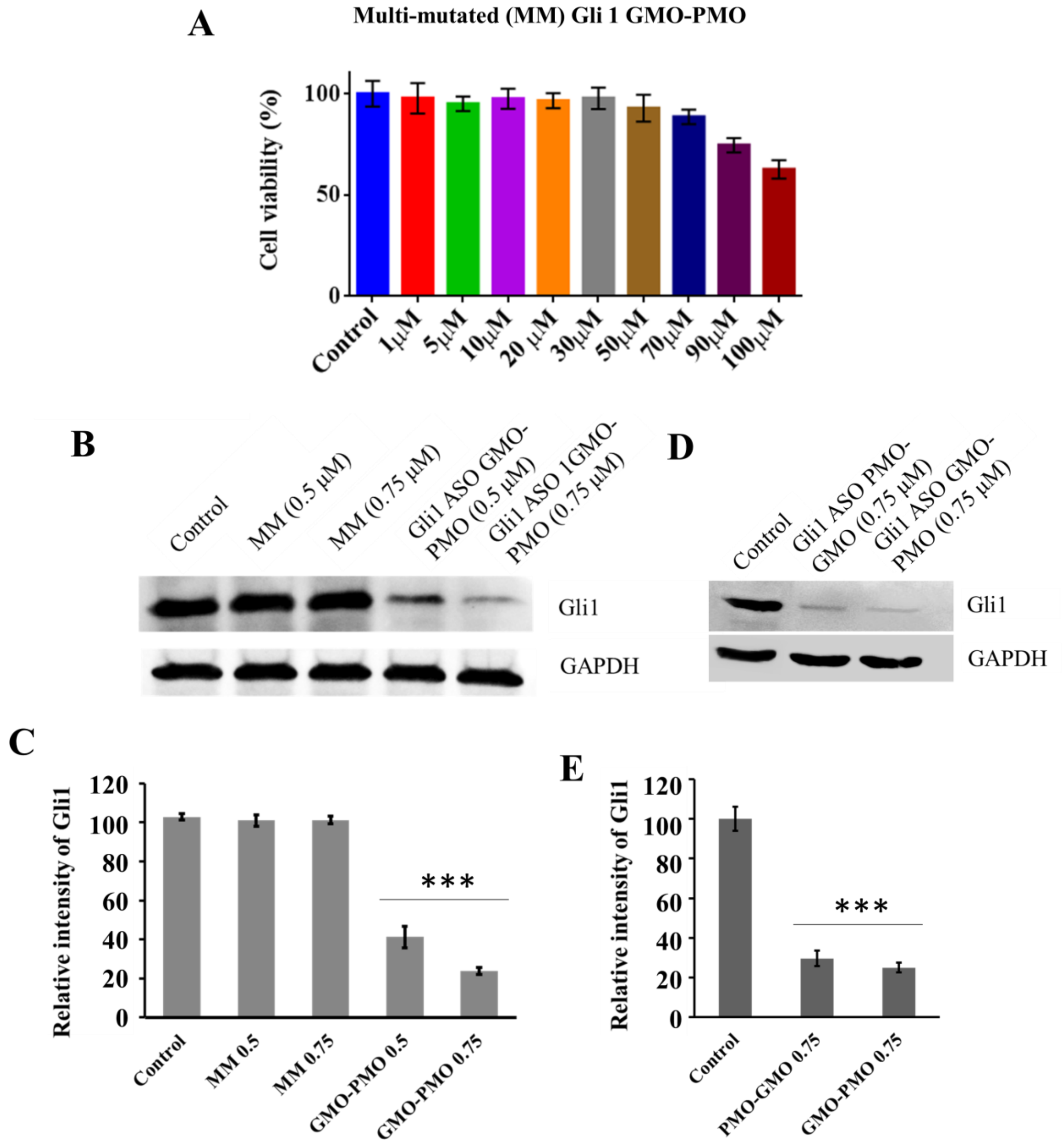


**Supporting Figure 2: Cytotoxicity and dose standardization of GMO PMO chimera. A** Cytotoxicity assay by MTT was performed in ShhL2 cells to evaluate the toxicity of GMO-PMO chimera. **B.** Standardization of dose of Gli1 ASO GMO-PMO for silencing the Gli1 protein expression in ShhL2 cells. **C.** Densitometry analysis of immunoblot data. **D.** Comparison of the efficacy of GMO-PMO and PMO-GMO chimera in ShhL2 cells. **E.** Densitometry analysis of immunoblot data.

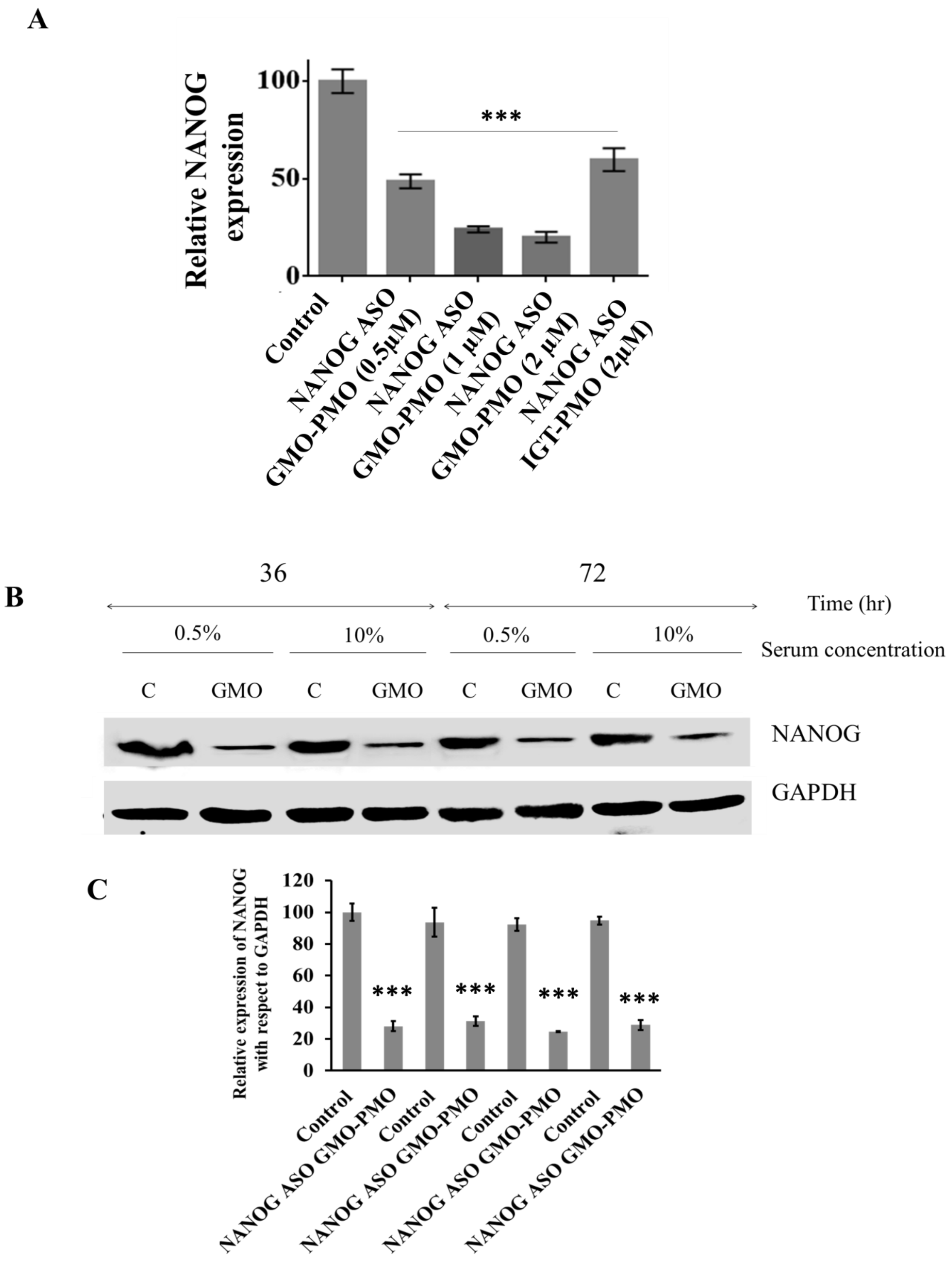


**Supporting Figure 3.** The densitometric analysis of immunoblot data of dose dependent inhibition of NANOG by NANOG ASO GMO-PMO. **B.** Immunoblot data shows the serum and time dependent inhibition of NANOG with NANOG ASO GMO-PMO. **C.** The densitometric analysis of immunoblot data.

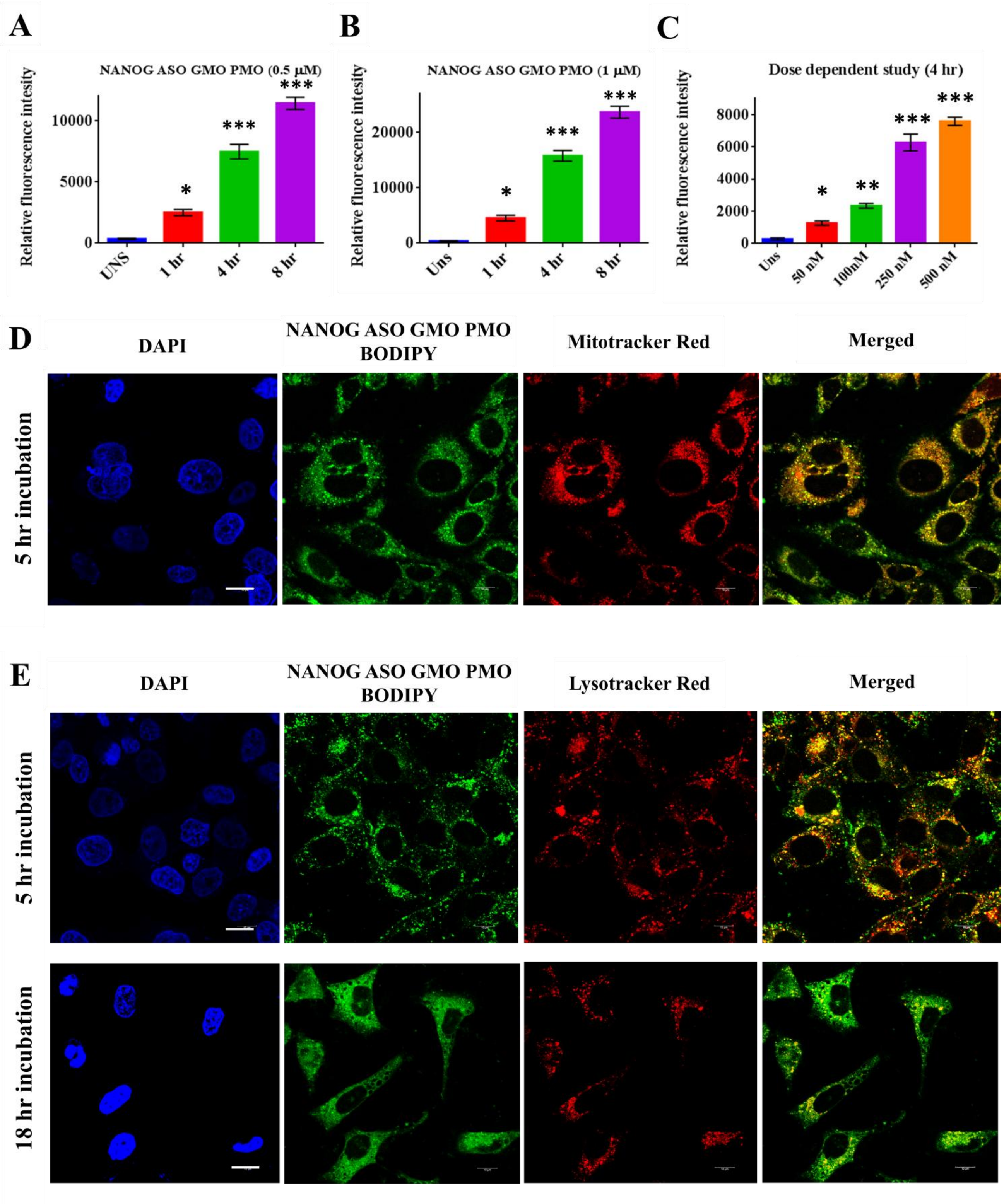


**Supporting Figure 4:** The bar diagrams show the time dependent entry of **A.** bodipy NANOG ASO GMO PMO (0.5 µM) and **B.** NANOG ASO GMO-PMO (1 µM). **C.** The bar diagram shows the dose dependent entry of compound in MCF-7 cells. **D.** The confocal images show the localization of NANOG ASO GMO-PMO in mitochondria. Nuclei were stained by DAPI and Mitotracker red was used to stain mitochondria. **E.** The images show the localization of antisense oligo in lysosome and their endosomal escape. Lysotracker red was used to stain lysosome. Error bars indicate means ± SE (n = 3), and data are presented as percentages relative to the non-treated MCF7 cells. *p< 0.05.

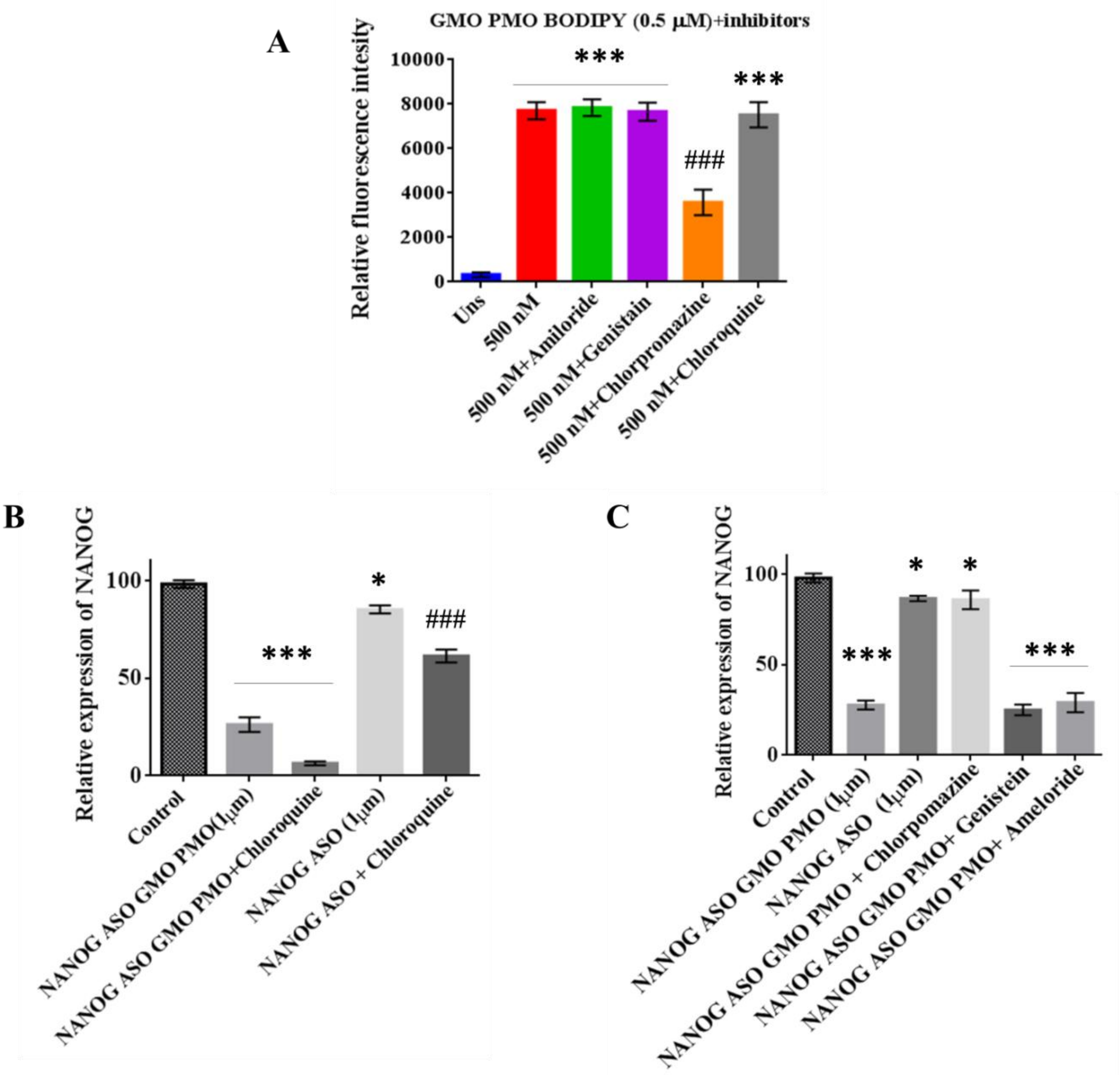


**Supporting Figure 5: A.** The bar diagrams show the entry of bodipy NANOG ASO GMO PMO (0.5 μM) in MCF7 cells in presence or absence of different inhibitors of endocytosis pathway. **B.** The bar diagram shows the densitometric analysis of immunoblot data, where the cells treated with NANOG ASO and NANOG ASO GMO-PMO in presence or absence of Chloroquine. **C.** The bar diagram represents the densitometric analysis of immunoblot of endocytosis inhibitor treated cells. Error bars indicate means ± SE (n = 3), and data are presented as percentages relative to the control MCF7 cells. *p< 0.05. # indicates the significant difference between control and NANOG ASO+Chloroquine treated cells.

**Control**

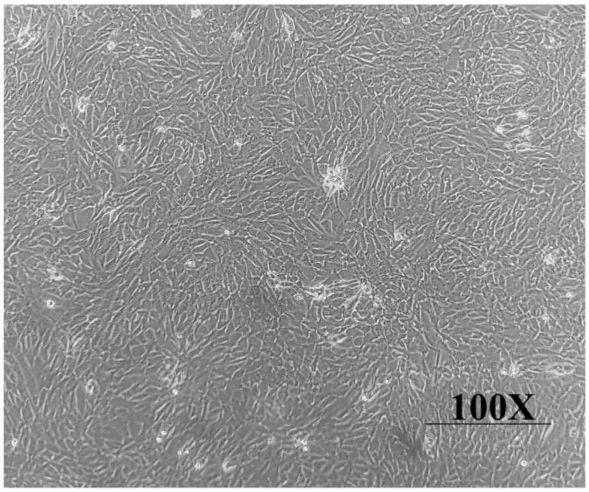


**NANOG ASO (1 µM)**

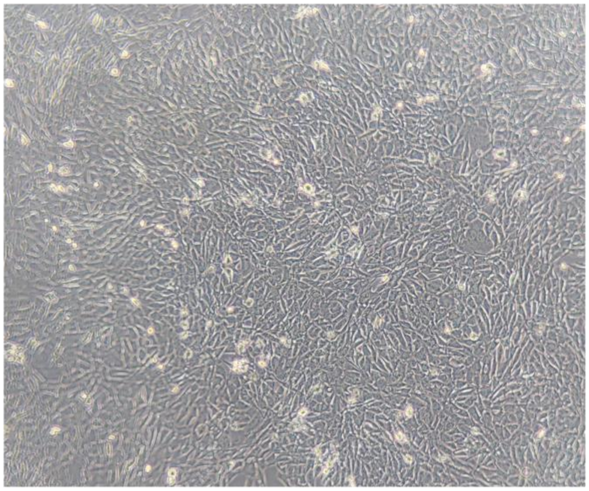

**NANOG ASO (1 µM)+Chloroquine**

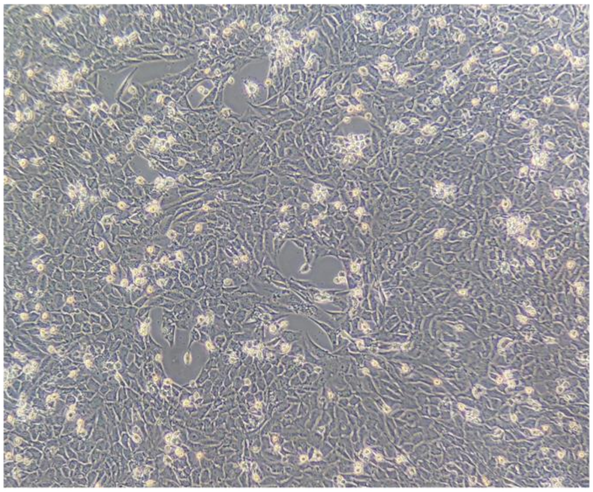

**NANOG ASO GMO-PMO (1 µM)**

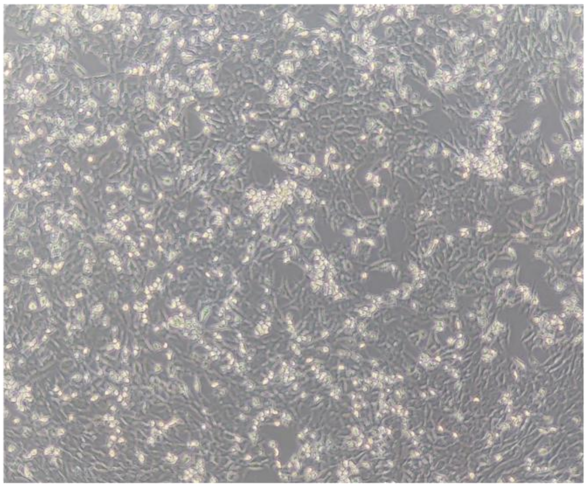

**NANOG ASO GMO-PMO (1 µM) +Chloroquine**

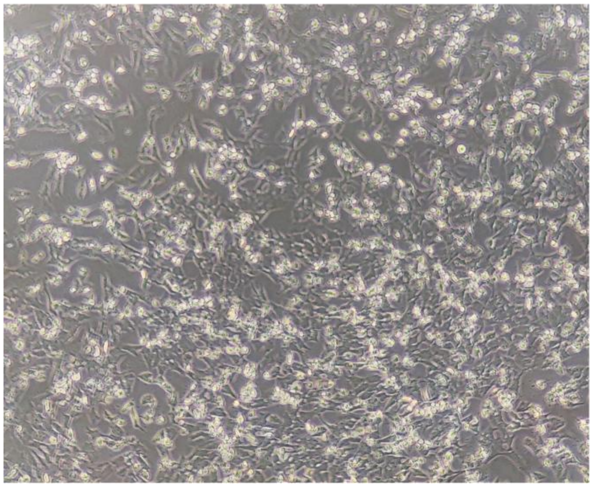

**NANOG ASO GMO-PMO + Chlorpromazine**

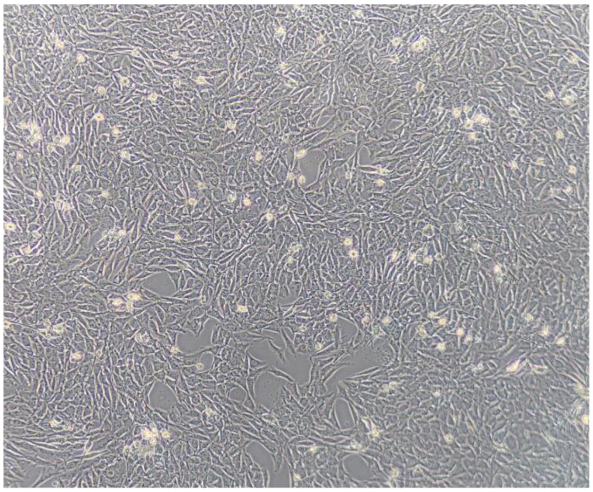

**NANOG ASO GMO-PMO + Genistein** | **NANOG ASO GMO-PMO + Amiloride**

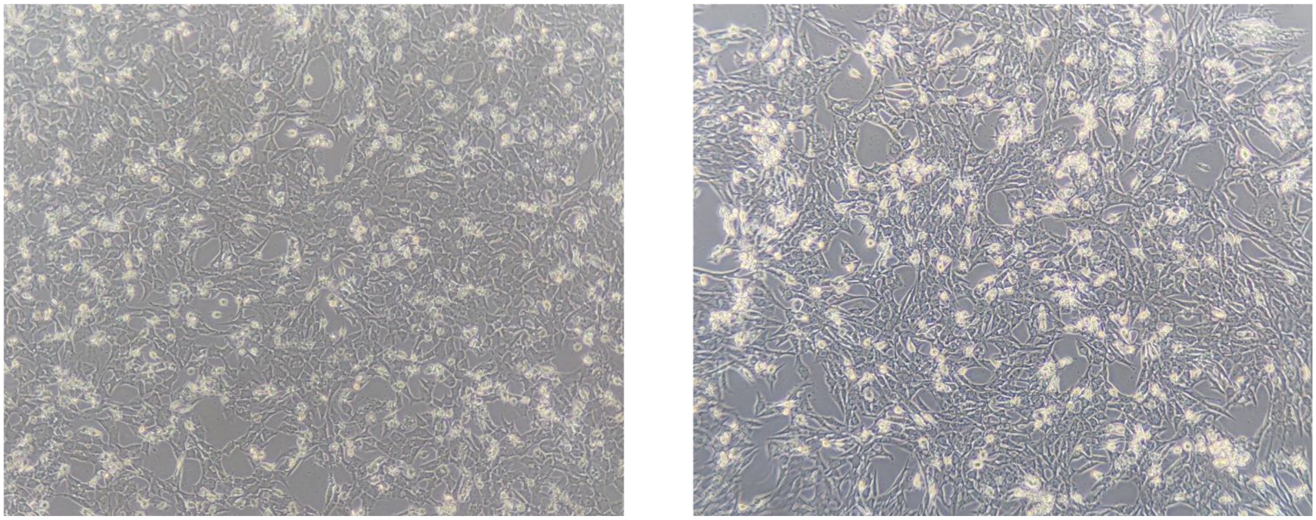

**Supporting Figure 6:** The morphological alteration in MCF7 cells after treatment with NANOG ASO, NANOG ASO in combination with Chloroquine, NANOG ASO GMO PMO, NANOG ASO GMO PMO in combination with Chloroquine, Chlorpromazine, Genistein and Ameloride.

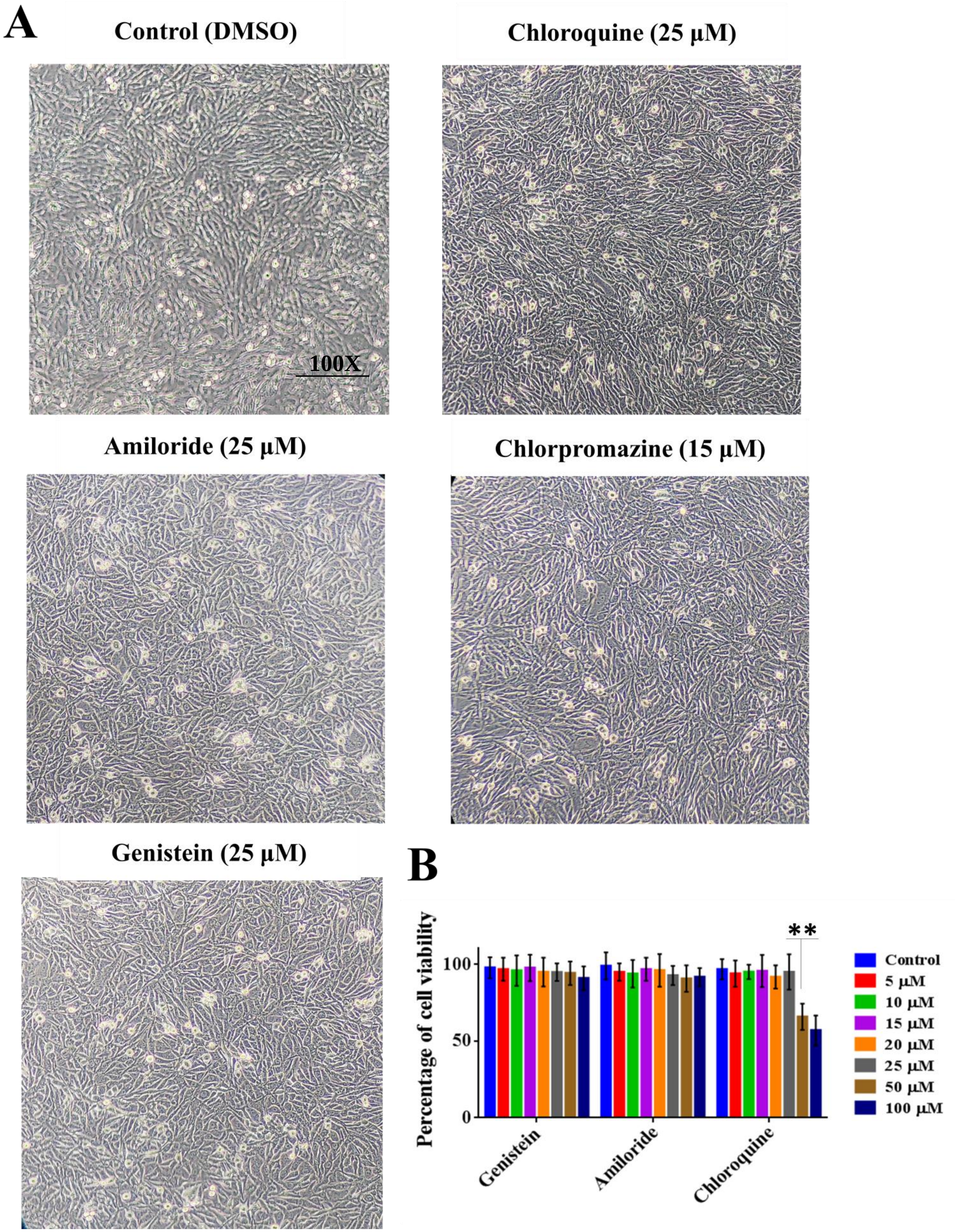
A
Control (DMSO)
100X
Chloroquine (25 µM)
Amiloride (25 µM)
Chlorpromazine (15 µM)
Genistein (25 µM)
B
Percentage of cell viability
0
50
100
**
Genistein
Amiloride
Chloroquine
Control
5 µM
10 µM
15 µM
20 µM
25 µM
50 µM
100 µM

**Supporting Figure 7: A.** The morphological alteration in MCF7 cells after treatment with Chloroquine, Chlorpromazine, Genistein and Ameloride alone. **B.** The MTT data shows the cytotoxicity of Chloroquine, Genistein and Ameloride. Error bars indicate means ± SE (n = 3), and data are presented as percentages relative to the control MCF7 cells. *p< 0.05.

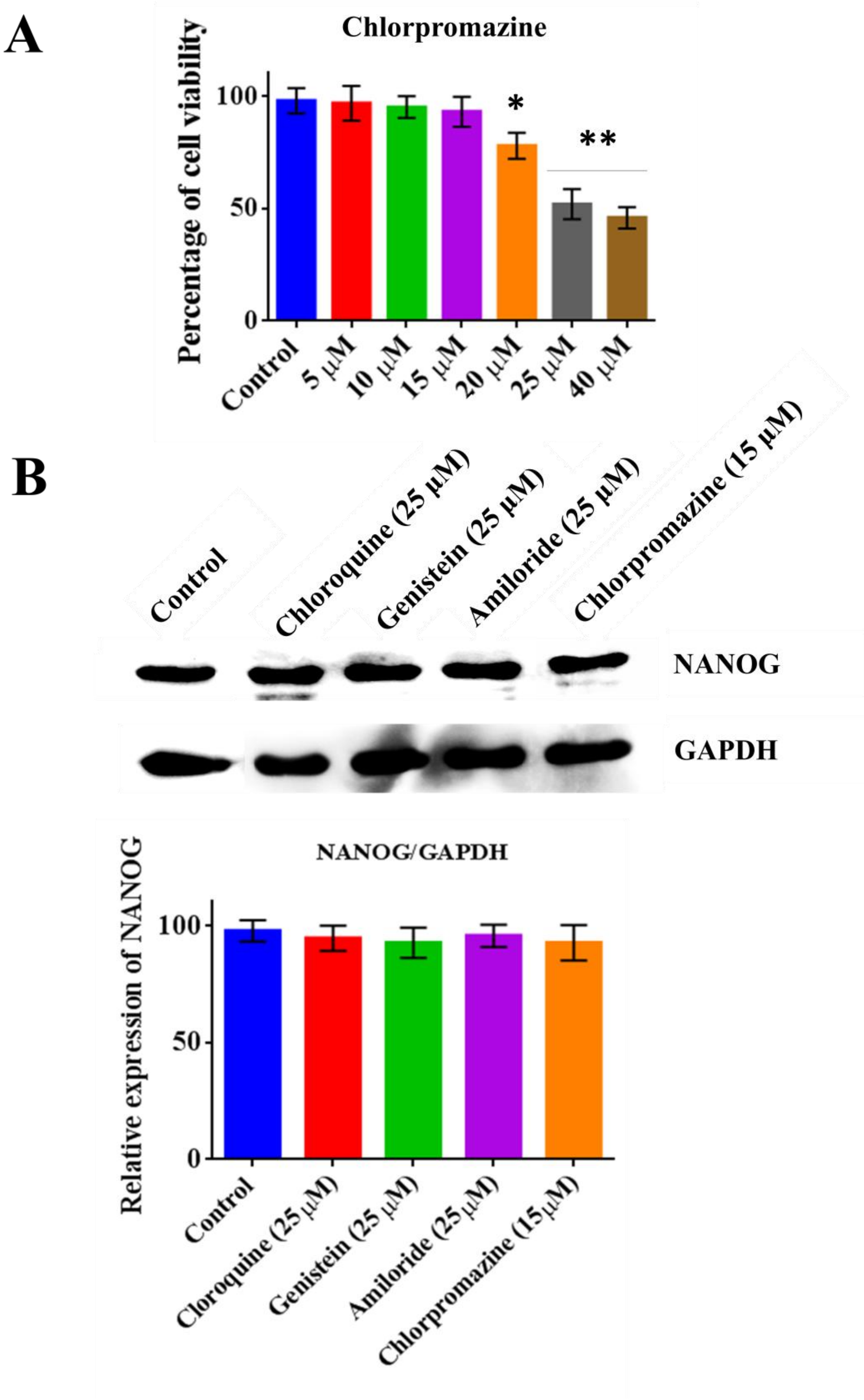

**Supporting Figure 8:** **A.** The MTT data shows the cytotoxicity of Chlorpromazine, **B.** The immunoblot data show the expression of NANOG it's densitometric analysis after treatment with the inhibitors alone. Error bars indicate means ± SE (n = 3), and data are presented as percentages relative to the control MCF-7 cells. *p< 0.05.

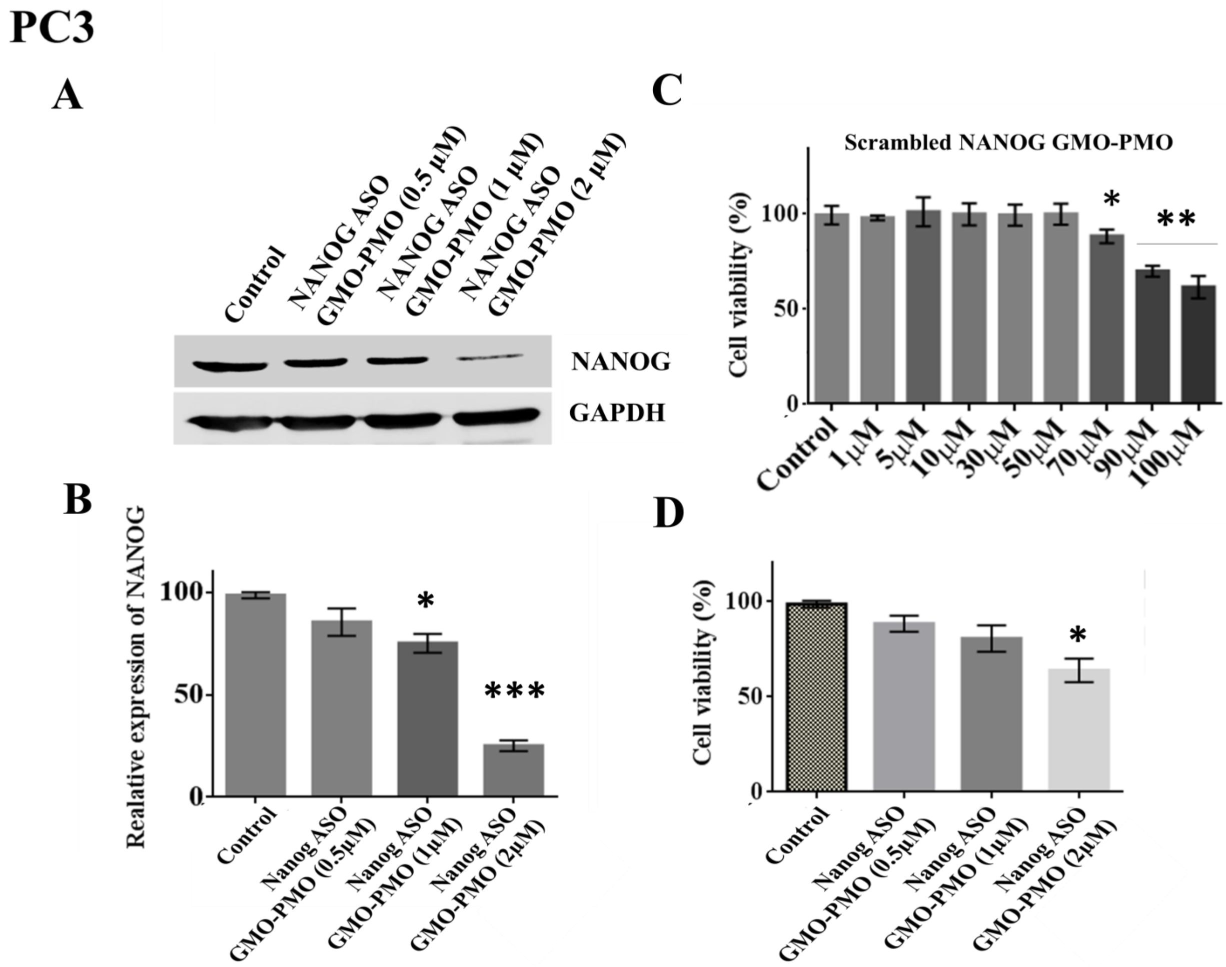


**Supporting Figure 9**: **A.** Immunoblot analysis and **B.** its graphical representation of the expression of NANOG in PC3 cells at different doses of NANOG ASO GMO PMO treatment. The protein expression levels are expressed relative to GAPDH. **C.** Cell viability is measured by MTT and its graphical depiction in PC3 cells after treatment with increasing doses of Scrambled NANOG GMO PMO and **D.** with different doses of NANOG ASO GMO PMO. Error bars indicate means ± SE (n = 3), and data are presented as percentages relative to the non-treated PC3 cells. *p< 0.05.

**PC3**

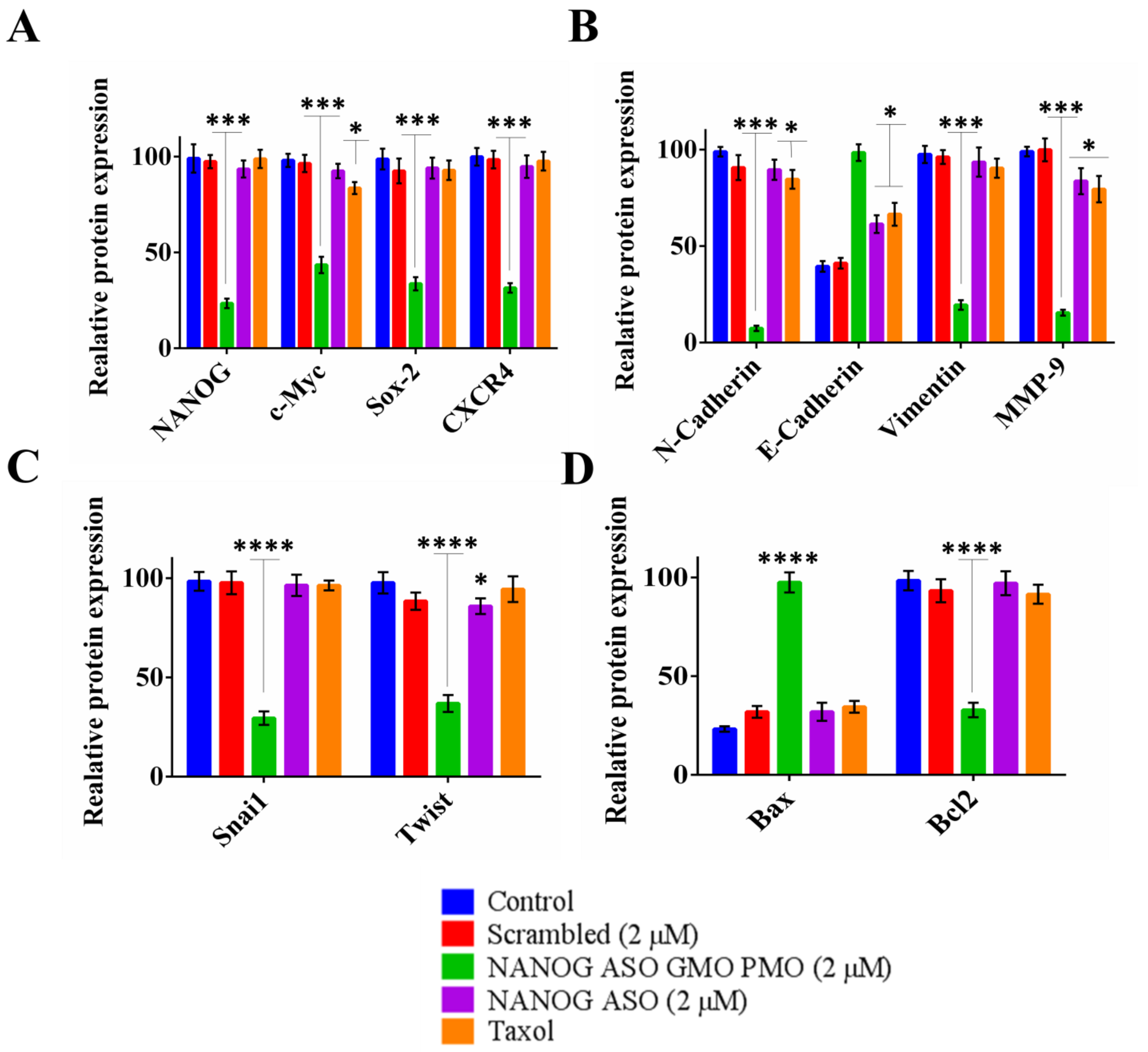


**Supporting Figure 10**: The densitometric analysis of expression of **A.** NANOG, c-Myc, Sox-2, CXCR4, **B.** N- Cadherin, E- Cadherin, Vimentin, MMP9, **C.** Snai1, Twist, and **D.** Bax, Bcl2 relative to GAPDH in PC3 cells. Error bars indicate means ± SE (n = 3), and data are presented as percentages relative to the non-treated PC3 cells. *p< 0.05.

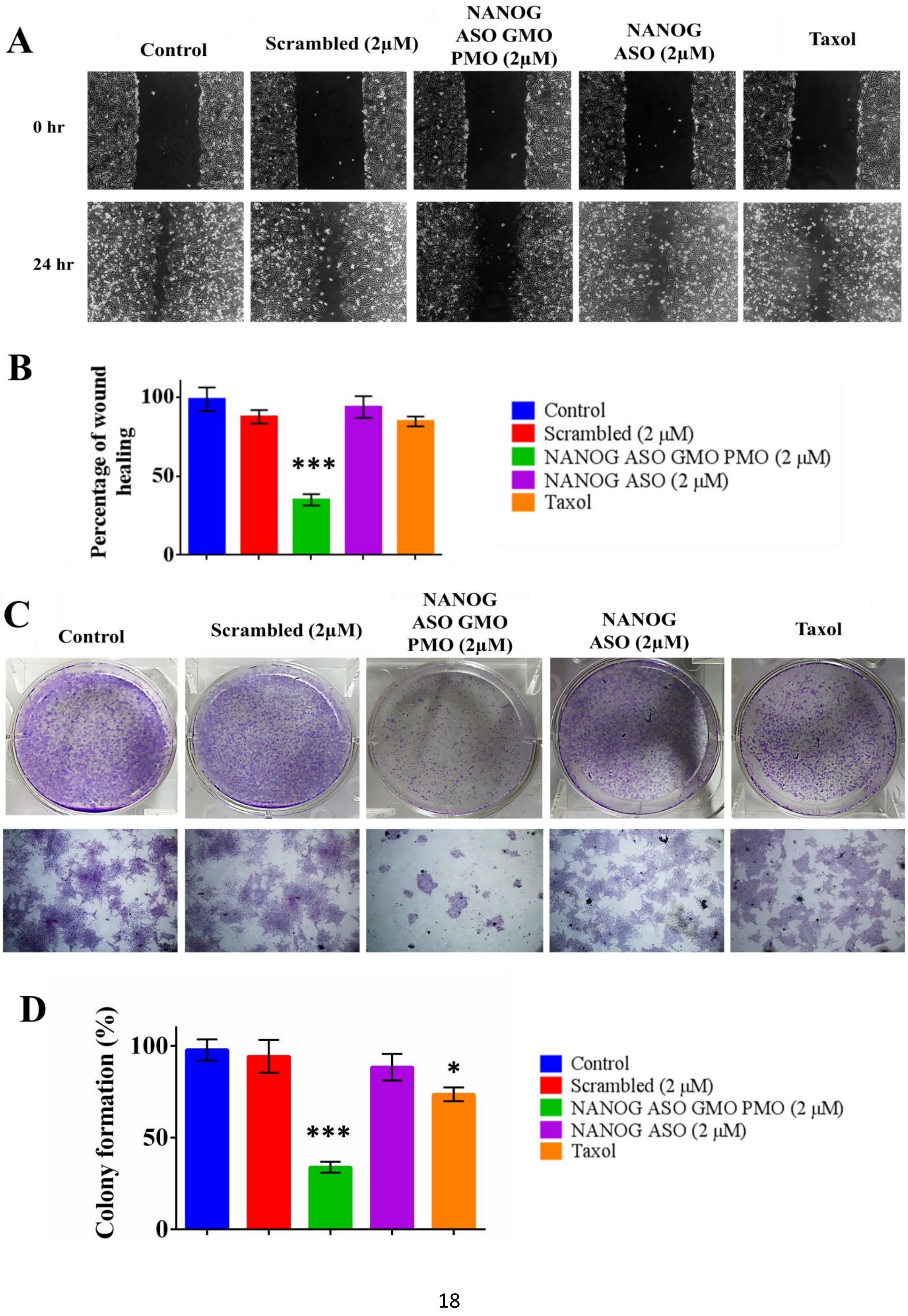

A
Control
Scrambled (2μM)
NANOG ASO GMO PMO (2μM)
NANOG ASO (2μM)
Taxol
0 hr
24 hr
B
Percentage of wound healing
100
50
0
***
Control
Scrambled (2 μM)
NANOG ASO GMO PMO (2 μM)
NANOG ASO (2 μM)
Taxol
C
Control
Scrambled (2μM)
NANOG ASO GMO PMO (2μM)
NANOG ASO (2μM)
Taxol
D
Colony formation (%)
100
50
0
***
*
Control
Scrambled (2 μM)
NANOG ASO GMO PMO (2 μM)
NANOG ASO (2 μM)
Taxol

**Supporting Figure 11**:**A.** Wound healing assay in the cells upon indicated treatment where the width of wound closure in PC3 cells at 0 h was set to 100%. **B.** Percentage of wound closure plotted graphically with regard to control. **C.** Anchorage-dependent colony formation assay. **D.** The percentage of cells forming colonies was plotted in bar diagram after different compound treatment conditions compared to that of parental PC3 cells. Scale bars represent 60 µm. Error bars indicate means ± SE (n = 3), and data are presented as percentages relative to the non-treated MCF-7 cells. *p< 0.05.

### MDA MB 231

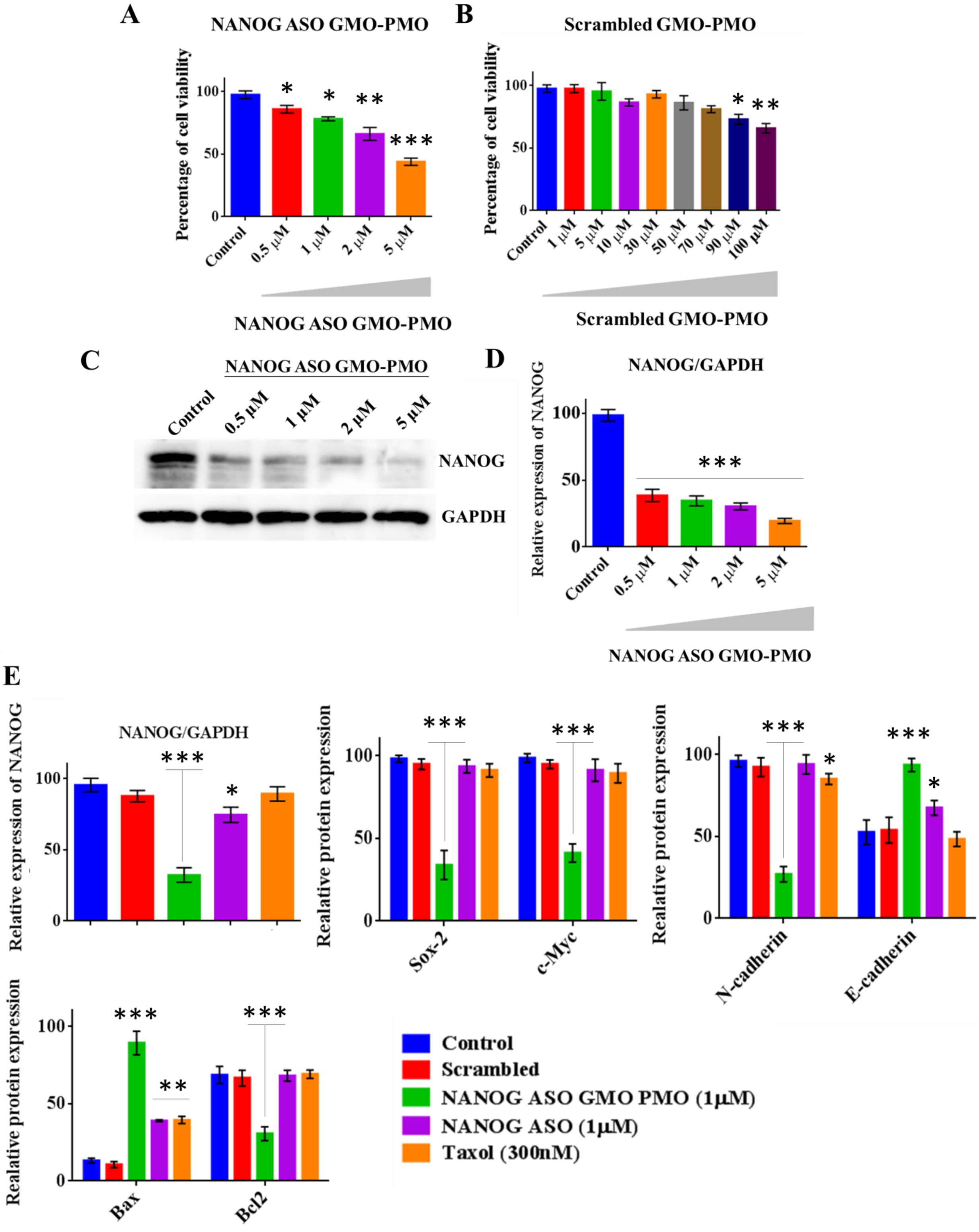
A
NANOG ASO GMO-PMO
Percentage of cell viability
Control
0.5 µM
1 µM
2 µM
5 µM
NANOG ASO GMO-PMO
B
Scrambled GMO-PMO
Percentage of cell viability
Control
1 µM
5 µM
10 µM
30 µM
50 µM
70 µM
90 µM
100 µM
Scrambled GMO-PMO
C
NANOG ASO GMO-PMO
Control
0.5 µM
1 µM
2 µM
5 µM
NANOG
GAPDH
D
NANOG/GAPDH
Relative expression of NANOG
Control
0.5 µM
1 µM
2 µM
5 µM
NANOG ASO GMO-PMO
E
NANOG/GAPDH
Relative expression of NANOG
Realative protein expression
Sox-2
c-Myc
Realative protein expression
N-cadherin
E-cadherin
Realative protein expression
Bax
Bcl2
Control
Scrambled
NANOG ASO GMO PMO (1µM)
NANOG ASO (1µM)
Taxol (300nM)

**Supporting Figure 12**: **A.** Cell viability is measured by MTT and its graphical depiction in MDA MB-231 cells after treatment with increasing doses of NANOG ASO GMO PMO and **B.** with different doses of Scrambled NANOG GMO PMO. **C.** Immunoblot analysis and **D.** its graphical representation of the expression of NANOG in MDA MB-231 cells at different doses of NANOG ASO GMO PMO treatment. **E.** Densitometric analysis of NANOG, c-Myc, Sox-2, N- Cadherin, E- Cadherin, Bax, Bcl2 immunoblot data relative to GAPDH in MDA MB-231 cells. The protein expression levels are expressed relative to GAPDH. Error bars indicate means ± SE (n = 3), and data are presented as percentages relative to the non-treated MDA MB-231 cells. *p< 0.05.

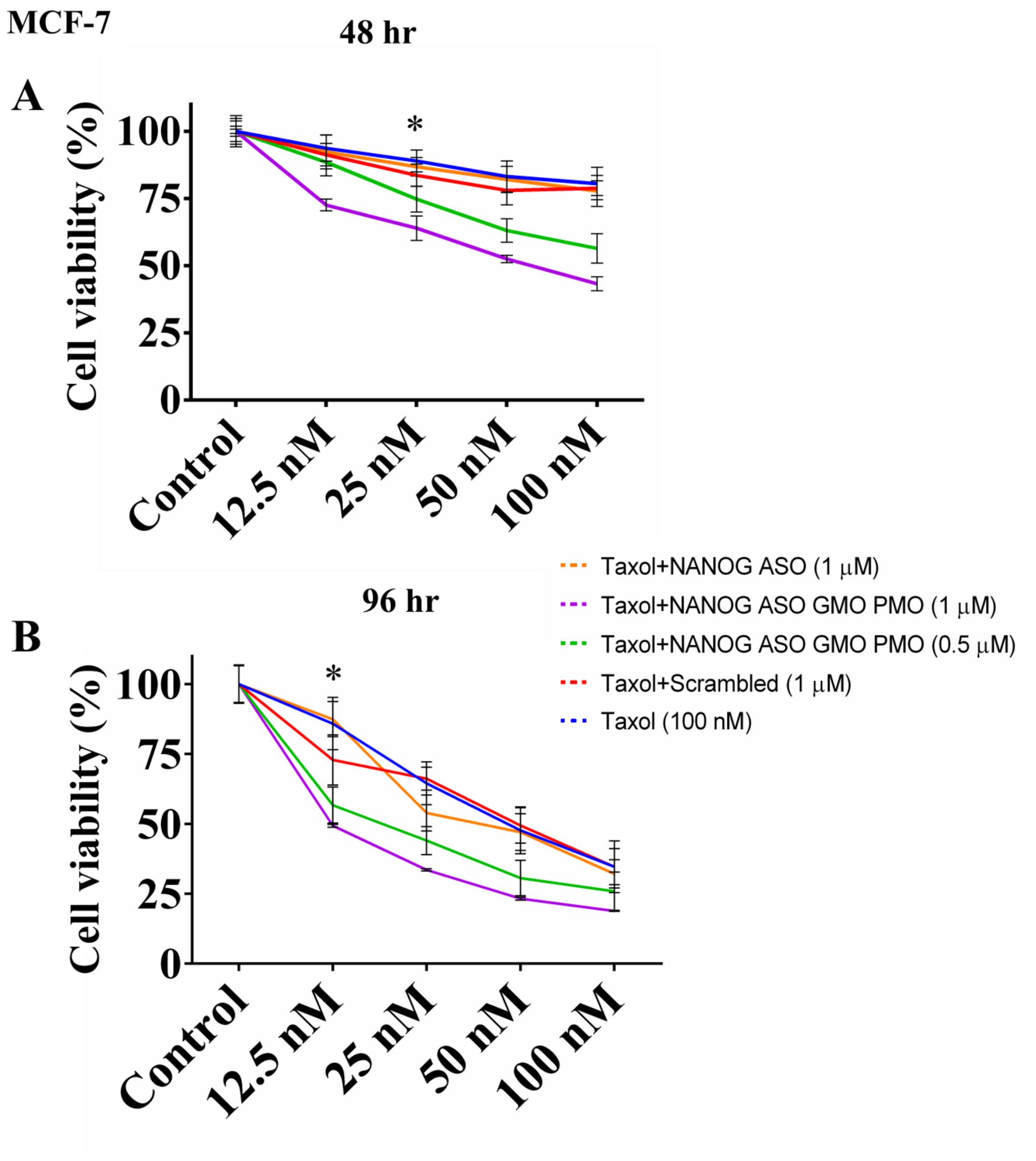

**Supporting Figure 13**: **A.** Graphical depiction of cell viability as measured by MTT where different oligo [Scrambled NANOG GMO PMO (1 µM), NANOG ASO (1 µM), NANOG ASO GMO PMO (0.5 µM) and NANOG ASO GMO PMO (1 µM)] treated MCF7 cells were subjected to 12.5nM, 25nM, 50nM and 100nM doses of taxol treatment for 48 and **B.** 96 hr respectively. Error bars indicate means ± SE (n = 3), and data are presented as percentages relative to the non-treated MCF-7 cells. *p< 0.05.

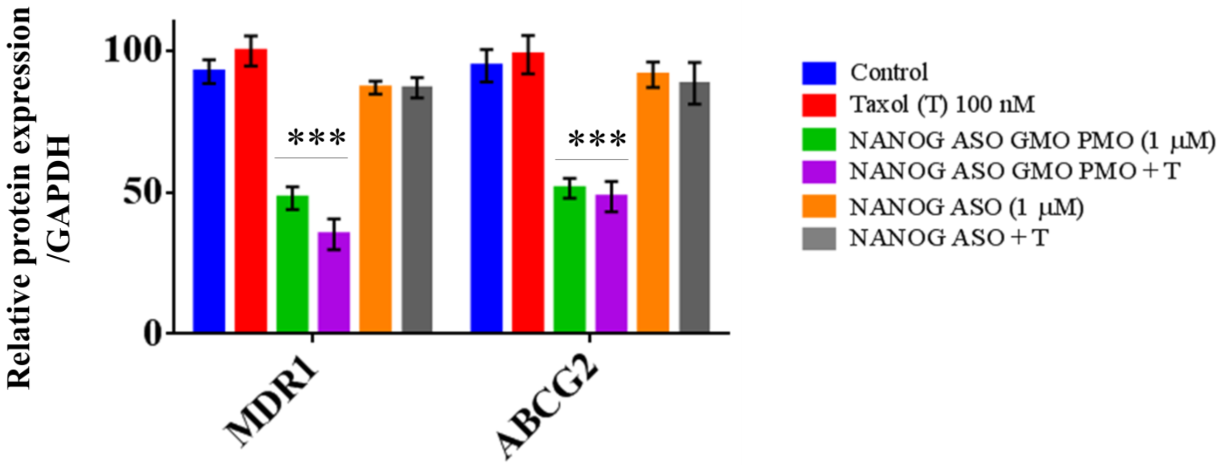


**Supporting Figure 14**: The bar diagram represents the densitometric analysis of MDR1 and ABCG2 immunoblot data. Error bars indicate means ± SE (n = 3), and data are presented as percentages relative to the non-treated MCF-7 cells. *p< 0.05.

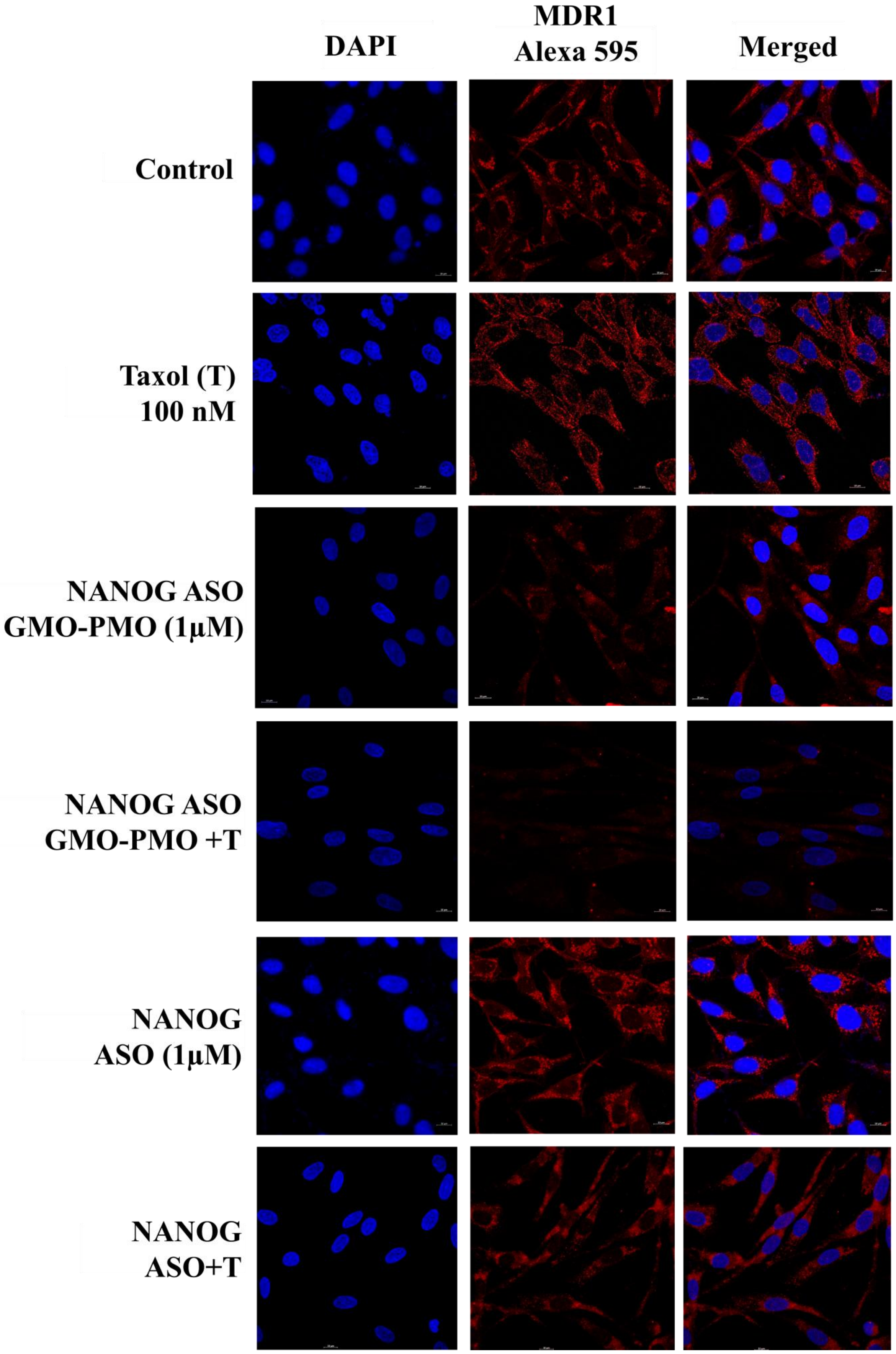
DAPI
MDR1
Alexa 595
Merged
Control
Taxol (T)
100 nM
NANOG ASO
GMO-PMO (1µM)
NANOG ASO
GMO-PMO +T
NANOG
ASO (1µM)
NANOG
ASO+T

**Supporting Figure 15**: Confocal immunofluorescence microscopic analysis of MDR1 (shown in red) protein in control and treated cells. Nuclei were stained with hoechst (blue).

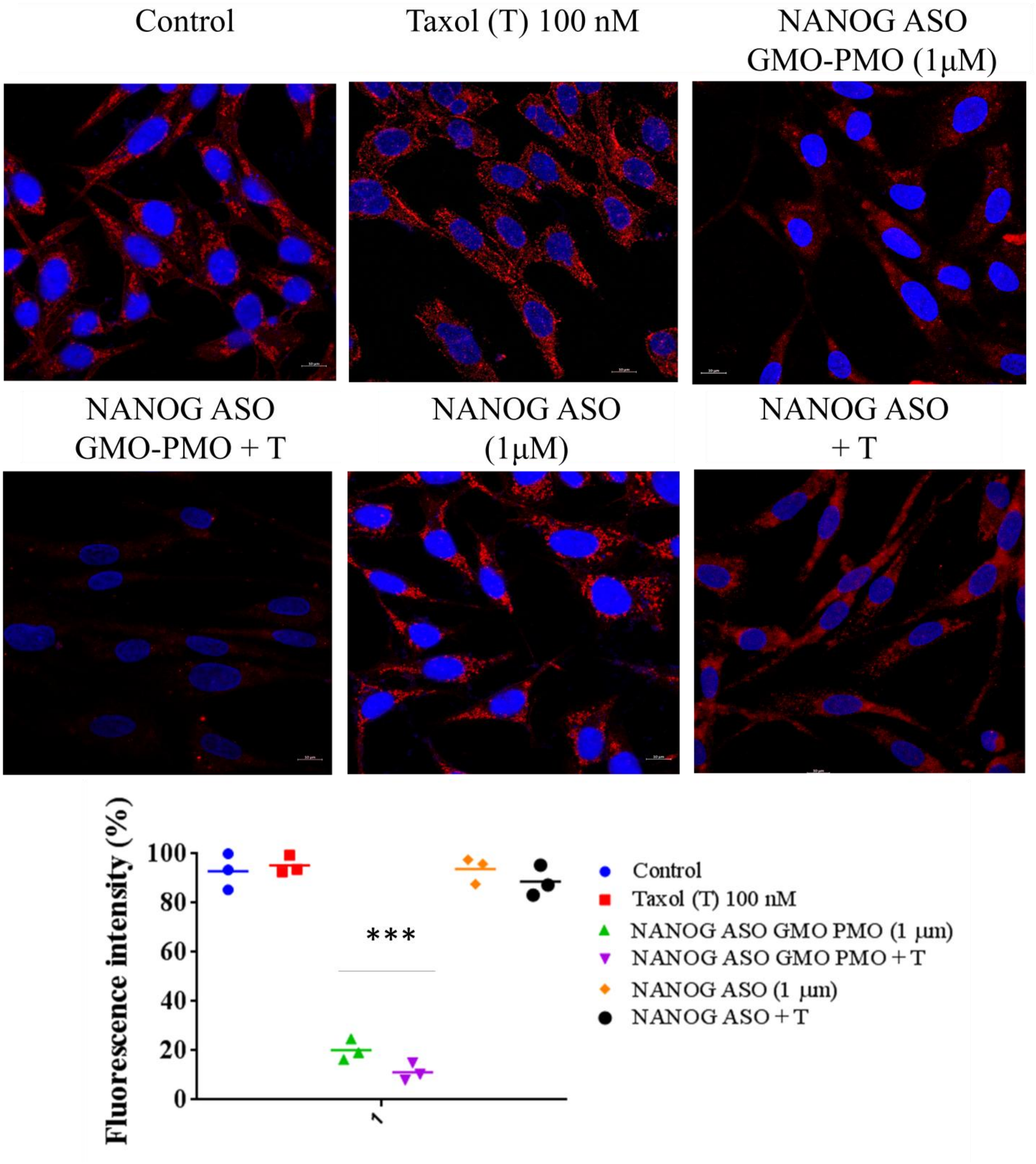


**Supporting Figure 16:** Enlarged view of merged images of supporting figure 15. Quantification of MDR1 intensity per nucleus was obtained from confocal immunofluorescence microscopy and was calculated for 20–25 cells and the values are represented as percentage. Error bars indicate means ± SE (n = 3), and data are presented as percentages relative to the non-treated MCF-7 cells. *p< 0.05.

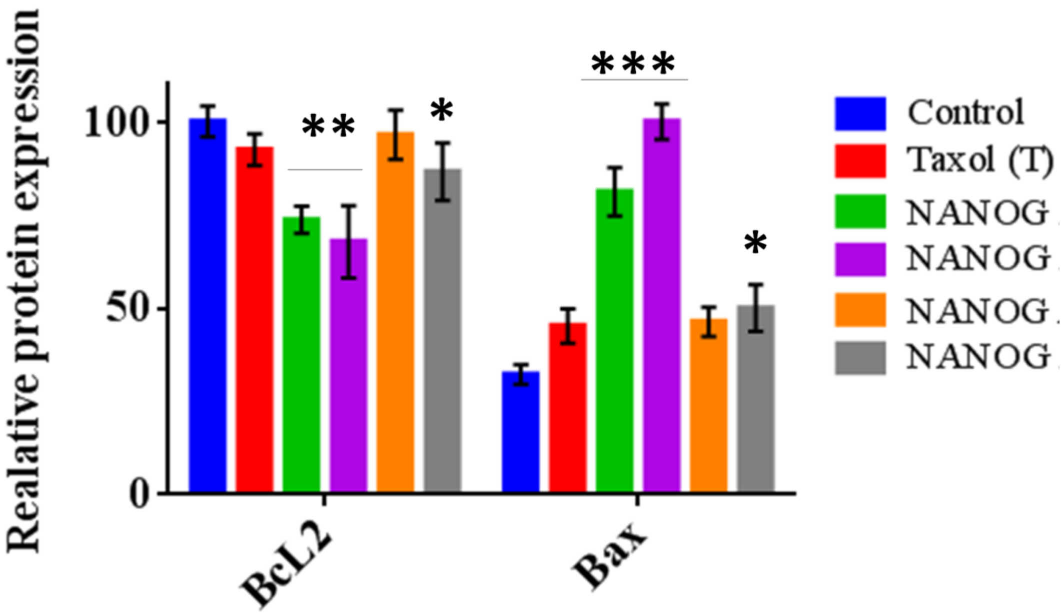


**Supporting Figure 17:** The densitometric analysis of Bax and Bcl2 immunoblot data. Error bars indicate means ± SE (n = 3), and data are presented as percentages relative to the non-treated MCF-7 cells. *p< 0.05.

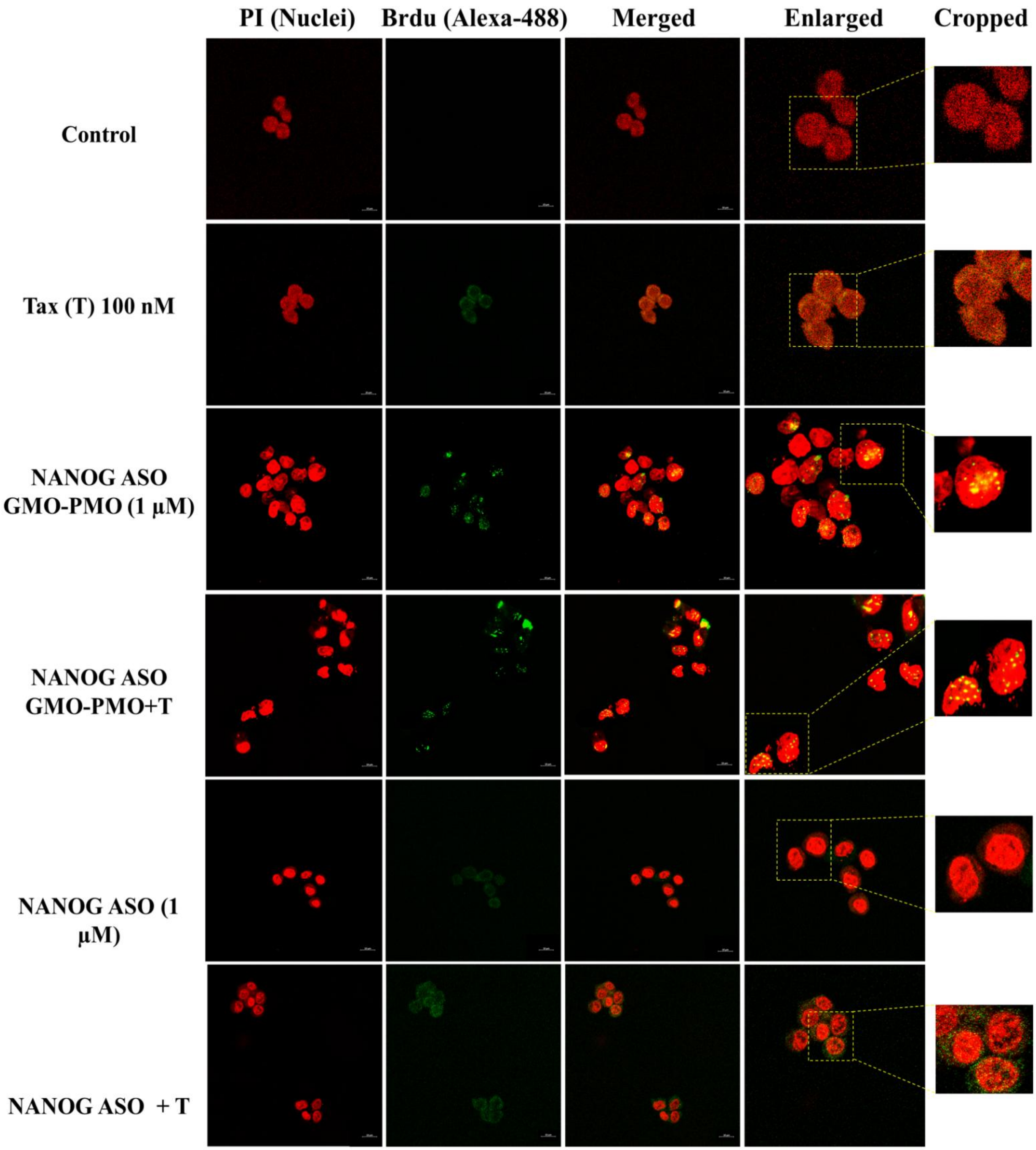
PI (Nuclei)
Brdu (Alexa-488)
Merged
Enlarged
Cropped
Control
Tax (T) 100 nM
NANOG ASO GMO-PMO (1 µM)
NANOG ASO GMO-PMO+T
NANOG ASO (1 µM)
NANOG ASO + T

**Supporting Figure 18:** The confocal images show the TUNEL formation by BrdU incorporation in MCF-7 cells. The nuclei were stained with PI and BrdU was stained by alexa 488 tagged anti BrdU antibody. The yellow dots on the merged images indicate the TUNEL formation or BrdU incorporation.

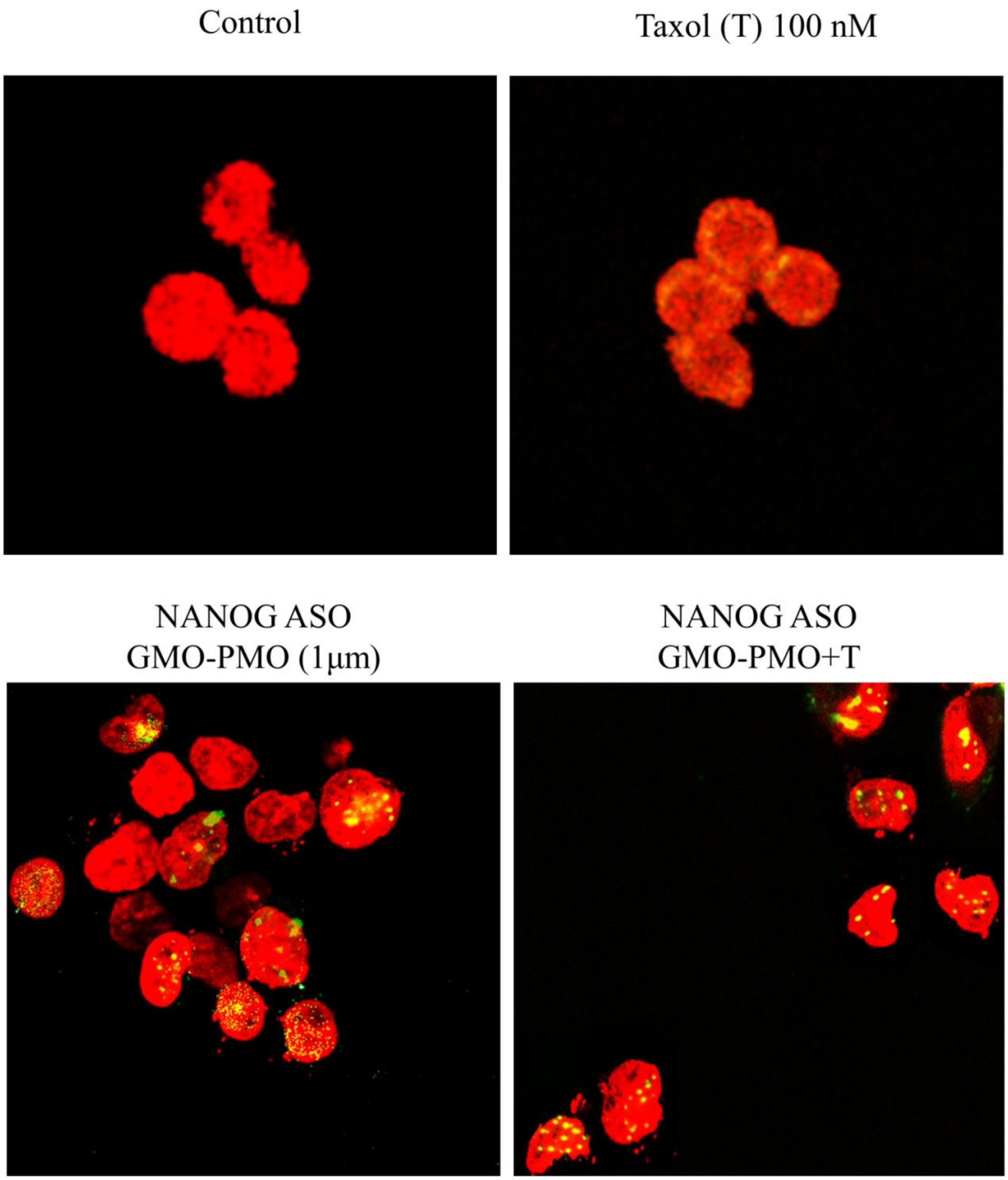


**Supporting Figure 19:** The enlarged merged images of supporting figure 18.

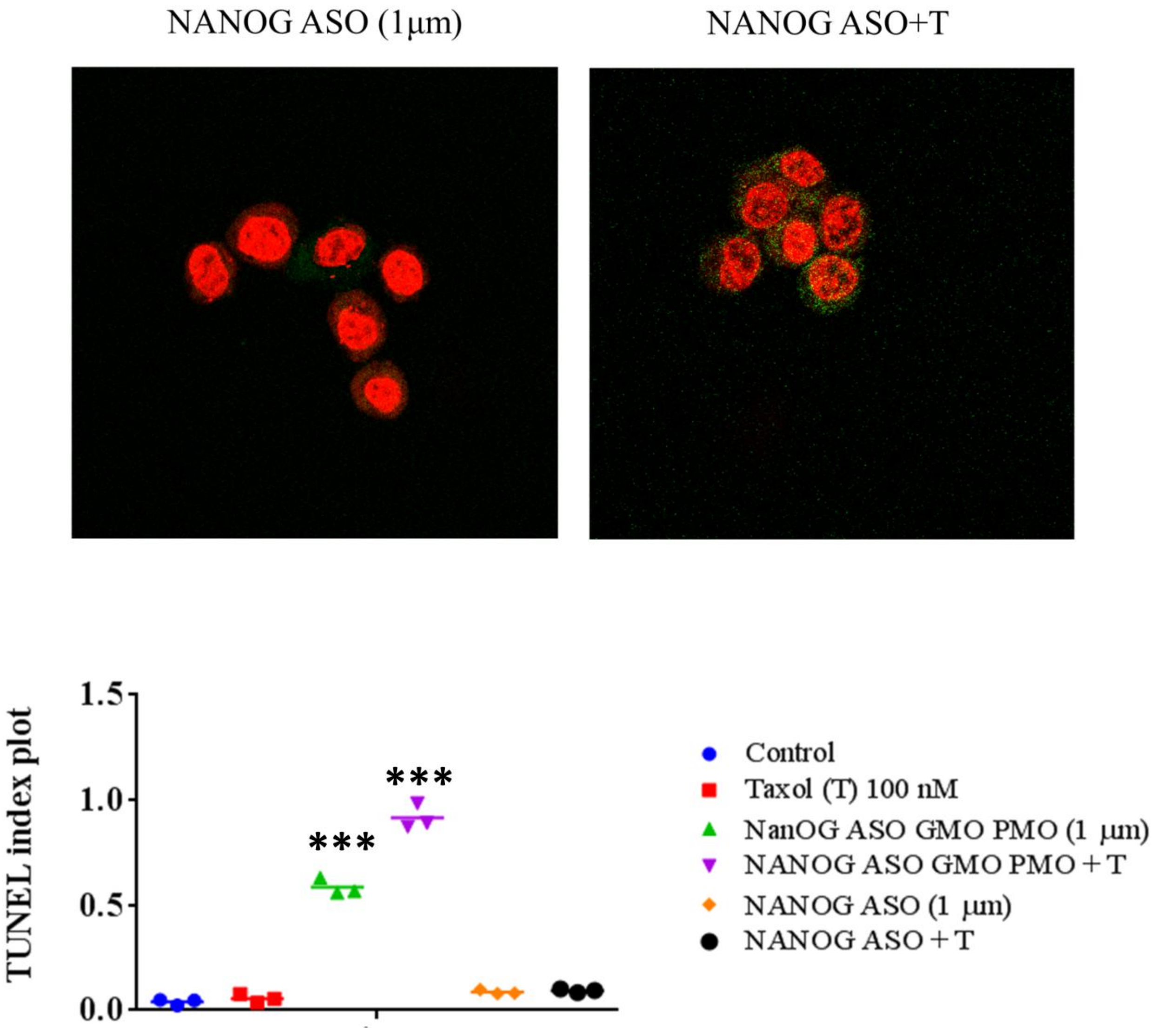


**Supporting Figure 20:** The enlarged merged images of supporting figure 18. Quantification of BrdU intensity per nucleus was obtained from confocal immunofluorescence microscopy and was calculated for 20–25 cells. Error bars indicate means ± SE (n = 3), and data are presented as number of TUNEL positive cells/total number of cells. *$p < 0.05$.
